# Accelerometer-derived classifiers for early detection of degenerative joint disease in cats

**DOI:** 10.1101/2024.12.13.628330

**Authors:** A.X. Montout, E Maniaki, T. Burghardt, M. J. Hezzell, E. Blackwell, A.W. Dowsey

## Abstract

Decreased mobility is a clinical sign of degenerative joint disease (DJD) in cats, which is highly prevalent, with 61% of cats aged six years or older showing radiographic evidence of DJD. Radiographs can reveal morphological changes and assess joint degeneration, but they cannot determine the extent of pain experienced by cats. Additionally, there is no universal objective assessment method for DJD-associated pain in cats. Developing an accurate evaluation model could enable earlier treatment, slow disease progression, and improve cats’ well-being.

This study aimed to predict early signs of DJD in cats using accelerometers and machine learning techniques. Cats were restricted to indoors or limited outdoor access, including being walked on a lead or allowed into enclosed areas for short periods. Fifty-six cats were fitted with collar-mounted sensors that collected accelerometry data over 14 days, with data from 51 cats included in the analysis. Cat owners assessed their cats’ mobility and assigned condition scores, validated through clinical orthopaedic examinations. The study group comprised 24 healthy cats (no owner-reported mobility changes) and 27 unhealthy cats (owner-reported mobility changes, suggestive of early DJD). Data were segmented into 60-second windows centred around peaks of high activity. Using a Support Vector Machine (SVM) algorithm, the model achieved 78% (confidence interval: 0.65, 0.88) area under the curve (AUC), with 68% sensitivity (0.64, 0.77) at 75% specificity (0.68, 0.79).

These results demonstrate the potential of accelerometry and machine learning to aid early DJD diagnosis and improve management, offering significant advances in non-invasive diagnostic techniques for cats.

## Introduction

Degenerative joint disease (DJD), is a condition characterised by the progressive and irreversible deterioration of the articular cartilage within the synovial joints. This deterioration goes beyond the natural wear and tear associated with aging, leading to pain, impaired joint function, and poor quality of life. The condition results in articular cartilage degeneration and pathological changes in periarticular tissues, which cause low grade inflammation in the joint and reduced range of motion together with pain on movement. Known risk factors for DJD in cats include late neutering, obesity, outdoor access and trauma (<u>Maniaki et al, 2021</u>). It is estimated that most older cats have DJD, with 61% of cats aged 6 years or older showing radiographic evidence of osteoarthritis (Slingerland et al, 2011; Lascelles et al, 2010).

Decreased mobility is a highly prevalent clinical sign of DJD in cats (Slingerland et al., 2011). Although radiographic evaluation provides an accurate diagnosis and assessment of DJD, identifying pain responses in cats through clinical examination can often be challenging as it heavily relies on interpreting behavioural cues that may not consistently reflect underlying pain.

The subjective nature of pain perception in animals complicates the assessment, as cats may not exhibit overt pain behaviours consistently across different individuals or stages of disease progression. Furthermore, owner separation coupled with the physical examination location can result in clinically significant increases in perceived stress in cats and compromise vital sign assessments. Whenever possible, physical examinations and procedures should take place with the owner present and should be conducted with separation from unfamiliar cats to minimize stress and improve the accuracy of clinical evaluations (Griffin et al., 2021).

Moreover, there is no universally accepted, objective method for gauging the pain referrable to DJD in cats. Although more experienced assessors can accurately perceive pain in cats, the challenge intensifies as radiographic findings do not always consistently align with pain perception, making DJD less obvious to many owners. Presently, there are no fully validated subjective or objective assessment systems for the assessment of chronic DJD-associated pain in cats. Hence, identifying subtle changes in activity patterns represents a potential objective approach for detecting or measuring pain linked to DJD (Slingerland et al, 2011; Lascelles et al, 2010; Lascelles et al, 2012).

The development of an accurate evaluation model could allow earlier detection of DJD in cats, facilitating earlier treatment and potentially improving quality of life. This study aimed to predict early signs of DJD in indoor cats with the use of accelerometers and machine learning techniques. The study hypothesis was that the effect of DJD would be more pronounced during periods of intense activity, such as when the cat performs behaviours such as jumping or moving at high speed.

## Materials and Methods

### Study group

The study was approved by the University of Bristol Animal Welfare and Ethical Review Body (VIN/18/026; 9 August 2018) and the University of Bristol’s Health Sciences Faculty Research Ethics Committee (69041; 04/07/2018). Cats were recruited from those already enrolled in the ongoing Bristol Cats study and via study advertisements (University of Bristol Langford Veterinary School campus, local veterinary practices) and social media.

The Bristol Cats study, launched in May 2010, aimed to investigate the causes of common behaviour patterns and diseases in cats. 2200 kittens were recruited through online applications via various social media channels, email or telephone. Owners were also contacted and encouraged to participate through registration processes facilitated by veterinary clinics, online platforms, and advertising in local communities. Participation involved completing questionnaires at specific intervals throughout the cat’s life: at ages approximately 8-16 weeks, 6-7 months, 12-13 months, 18-19 months, 2.5 years, 4 years, and annually thereafter, contingent on study funding. These questionnaires gathered data on various aspects of cat management, such as diet and lifestyle, and examined associations between these factors and behaviours or conditions like aggression or obesity. Ethical approval was obtained to ensure the welfare of participating cats, with measures in place to report suspected neglect. Confidentiality of owner data was strictly maintained, and participant consent was obtained for data storage and usage. The study’s findings are intended to inform future improvements in cat health and welfare, paralleling the impact of similar longitudinal studies in human health, such as the “Children of the 90s” cohort study (Boyd et al, 2012).

For this study cats aged six years or older, living entirely indoors or with restricted outdoor access, were included in the study if their owners lived within 100 miles of the University of Bristol Veterinary School and agreed to equip them with an activity monitor on their collars for two weeks. The exclusion criteria for the study were as follows; being fearful of strangers, unable to acclimatise to wearing a collar (if not already wearing one) or unduly stressed by the process, receiving dietary supplements if initiated <30 days before the visit or if initiated >30 days before the visit and not continued during the study, receiving anti-inflammatory or analgesic medications, or having been diagnosed with any condition that could influence mobility (Maniaki et al, 2023).

Cats were assigned to case or control groups based on their mobility score (MS), derived from a subset of 12 mobility-related questions selected from the Bristol Cats questionnaire (see supplementary material). These questions were chosen for their relevance to DJD. Control cats were identified by an MS of 0, indicating no owner-assessed mobility impairment, whereas cats with an MS greater than 1 were assigned to the case group, suggesting owner-assessed mobility impairment most likely due to DJD rather than other disease processes. The MS calculation excluded scores of 1 to minimize uncertainty. Clinical examination findings and questionnaire data, including the Functional Mobility Profile Index (FMPI) (Benito et al, 2013) further informed group assignment. VetMetrica (Davies et al, 2020) questionnaires were also conducted, which were used to assess various aspects of a cat’s health and well-being, including chronic pain, quality of life, and appetite. For cats that also took part in the Bristol Cats study their last completed questionnaire where used, whereas the other completed those 12 questions online.

A clinical examination was conducted at the cats’ homes by a qualified veterinary surgeon, evaluating body condition score, orthopaedic health, joint pain, and temperament. The assessor remained unaware of each cat’s MS and questionnaire responses while performing the orthopaedic examination subsequently, activity monitors were attached to the cats’ collars for two weeks, the cats that were not wearing collars before were habituated before equipping them with a monitor (Maniaki et al, 2023).

For this study, Actical ® Z wearable devices (Philips Respironics) were used. The Actical activity monitor, weighing 16g, measures acceleration changes with a piezoelectric accelerometer. The device was set to output activity counts every 1 second. The raw data takes the form of MS Excel spreadsheets that contain the sensor data, in addition to a metadata spreadsheet (Table 1 A,B,C) which describes the mobility score as well as other metrics such as the age (Fig. S1) and the sex of the cats. To avoid visit-related effects, monitoring began the morning after placement. Owners maintained diaries noting any device or collar detachment. The study length was 14 days, and each cat was assigned to the DJD or control group before the start of accelerometer data acquisition; once assigned, cats remained in their designated groups for the duration of the study.

**Table 1A.** Metadata available for the cat population of the study included the cats excluded for analysis (40, 600, 655). Each cat was assigned a mobility score assessed by its owner. The Status was given based on the mobility score, for a score > 1 the status is 1 otherwise it is 0. The age (in years) of the cat at the start of the study was also reported.

| Cat ID | Breed | Pure Breed | Sex* | Age | Outdoor* | Status* | Mobility score | Body condition score |
| --- | --- | --- | --- | --- | --- | --- | --- | --- |
| 11 | BSH | 0 | 0 | 7.8 | 0 | 0 | 0 | 6 |
| 12 | DSH | 1 | 1 | 7.8 | 0 | 0 | 0 | 5 |
| 13 | DSH | 1 | 1 | 7 | 0 | 1 | 4 | 4 |
| 24 | Oriental | 0 | 0 | 7.8 | 0 | 1 | 4 | 6 |
| 26 | Asian | 1 | 0 | 7.8 | 0 | 0 | 0 | 5 |
| 36 | Ragamuffin | 0 | 0 | 8.3 | 0 | 1 | 4 | 7 |
| 40 | DSH | 1 | 0 | 8.4 | 0 | 0 | 0 | 7 |
| 57 | DSH | 1 | 1 | 6.8 | 1 | 0 | 0 | 5 |
| 75 | DSH | 1 | 1 | 6.8 | 0 | 0 | 0 | 5 |
| 77 | DSH | 1 | 0 | 7.7 | 0 | 0 | 0 | 7 |
| 78 | DSH | 1 | 1 | 6.8 | 0 | 0 | 0 | 7 |
| 128 | DSH | 1 | 0 | 7 | 1 | 0 | 0 | 6 |
| 129 | Ragdoll x | 1 | 1 | 7.2 | 0 | 0 | 0 | 7 |
| 600 | DSH | 1 | 1 | 11.5 | 0 | 1 | 7 | 6 |
| 601 | DSH | 1 | 0 | 6.5 | 0 | 0 | 0 | 5 |
| 602 | DSH | 1 | 1 | 6.7 | 0 | 0 | 0 | 5 |
| 603 | DSH | 1 | 1 | 20 | 0 | 1 | 8 | 4 |
| 604 | DSH | 1 | 0 | 10.4 | 0 | 0 | 0 | 6 |
| 605 | Siberian x | 1 | 1 | 11.6 | 0 | 1 | 4 | 4 |
| 607 | DSH | 1 | 0 | 15 | 0 | 1 | 2 | 7 |
| 608 | DSH | 1 | 1 | 10 | 0 | 1 | 4 | 6 |
| 609 | Bengal | 0 | 0 | 6.4 | 0 | 0 | 0 | 5 |
| 610 | Ragdoll x | 1 | 0 | 6.4 | 0 | 0 | 0 | 6 |
| 611 | Snowshoe | 0 | 1 | 9.4 | 1 | 0 | 0 | 4 |
| 612 | DSH | 1 | 0 | 8.7 | 0 | 0 | 0 | 5 |
| 613 | Burmese | 0 | 1 | 15.3 | 0 | 1 | 10 | 4 |
| 614 | Burmese | 0 | 1 | 13.1 | 0 | 1 | 6 | 6 |
| 618 | DSH | 1 | 0 | 10.8 | 0 | 1 | 4 | 7 |
| 619 | DSH | 1 | 1 | 11.1 | 0 | 1 | 7 | 6 |
| 620 | DSH | 1 | 0 | 8.2 | 0 | 0 | 0 | 5 |
| 621 | DSH | 1 | 1 | 9.2 | 0 | 0 | 0 | 6 |
| 622 | DSH | 1 | 1 | 11.1 | 0 | 1 | 4 | 6 |
| 626 | Devon Rex | 0 | 0 | 9 | 0 | 1 | 10 | 3 |
| 627 | DSH | 1 | 1 | 9.2 | 0 | 1 | 4 | 4 |
| 630 | Ragdoll | 0 | 1 | 11.3 | 0 | 1 | 3 | 5 |
| 631 | DSH | 1 | 1 | 6.3 | 0 | 1 | 4 | 4 |
| 632 | Bengal | 0 | 0 | 7.6 | 0 | 0 | 0 | 4 |
| 634 | Persian | 0 | 1 | 12 | 0 | 0 | 0 | 4 |
| 635 | DSH | 1 | 0 | 7.5 | 0 | 0 | 0 | 5 |
| 637 | DSH | 1 | 1 | 16.6 | 0 | 1 | 4 | 5 |
| 642 | BSH | 0 | 0 | 7.4 | 0 | 1 | 13 | 7 |
| 645 | DSH | 1 | 1 | 16.2 | 0 | 1 | 4 | 5 |
| 646 | DSH | 1 | 0 | 6.4 | 0 | 1 | 3 | 5 |
| 647 | DSH | 1 | 1 | 8.5 | 0 | 1 | 6 | 7 |
| 648 | DSH | 1 | 0 | 15 | 0 | 1 | 3 | 6 |
| 649 | DSH | 1 | 0 | 8.6 | 0 | 1 | 4 | 6 |
| 650 | DSH | 1 | 0 | 8.6 | 0 | 1 | 4 | 7 |
| 651 | Siamese x<br>Havana | 1 | 0 | 14.5 | 0 | 0 | 0 | 5 |
| 652 | Siamese x<br>Havana | 1 | 0 | 14.5 | 0 | 0 | 0 | 6 |
| 655 | DSH | 1 | 0 | 10.9 | 0 | 0 | 0 | 5 |
| 658 | DSH | 1 | 1 | 11.8 | 0 | 1 | 3 | 6 |
| 660 | Cornish<br>Rex x | 1 | 1 | 13.3 | 0 | 1 | 6 | 4 |
| 661 | Singapura x<br>Egyptian<br>Mau | 1 | 1 | 6.4 | 0 | 1 | 3 | 4 |
| 662 | Exotic<br>Short Hair | 0 | 0 | 14.1 | 0 | 1 | 4 | 4 |
| 11 | BSH | 0 | 0 | 7.8 | 0 | 0 | 0 | 6 |

\* In the “Status” column, 0 = DJD, 1 = no-DJD.
\* In the “Sex” column, 0 = Male, 1 = Female.
\*In the “Outdoor” column, 0 = indoors, 1 = limited outdoors access.
DSH: Domestic Short Hair
BSH: British Shorthair

**Table 1B.** Age and mobility score summary.

| **Health Status** | **Age Mean ± SD** | **Mobility Score**<br>**Median (Min, Max)** | **Body condition Score**<br>**Median (Min, Max)** |
| --- | --- | --- | --- |
| 0 (no-DJD) | 8.50 ± 2.26 | 1.00 (0.83, 1.00) | 5 (4, 7) |
| 1 (DJD) | 11.20 ± 3.54 | 0.83 (0.46, 1.00) | 6 (3, 7) |

**Table 1C.**
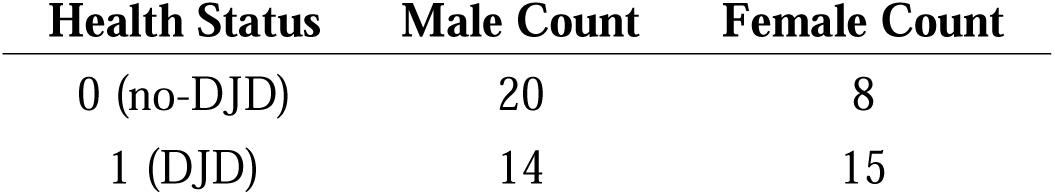
Sex distribution.

| **Health Status** | **Male Count** | **Female Count** |
| --- | --- | --- |
| 0 (no-DJD) | 20 | 8 |
| 1 (DJD) | 14 | 15 |

### Data acquisition

A total of four sensors were utilised across all the cats, with each sensor being passed on to another cat upon completion of data collection. For five cats, the sensors were configured to measure activity counts at a 33Hz sampling frequency, while for the remaining cats, the sensors recorded activity counts every second. The monitors configured to record accelerometery data at 33Hz files were too large (more than 1Gb), which made downloading the data from the device take up to 12 hours because of the device limited download bandwidth, this prevented quick turnover to use the accelerometer for the next cat, therefore the accelerometers were set to record at 1Hz making data files more manageable. All the activity data were therefore resampled to the second resolution (i.e. each count contains the sum of activity counts within each second) if it was not already at that resolution.

In Fig. S2 we show the activity data of all the cats including those not used in the analysis (3 cats). Each row shows the activity count data of a single cat throughout the entirety of the study.

### Building samples for machine learning

A supervised machine learning (ML) approach was used, which requires samples to be built that are composed of an array of features (activity counts) and a ground truth (outcome) value i.e. health score. To overcome the limitations of a relatively small sample size the dataset was augmented using a combinatorial approach.

To reflect the study hypothesis that DJD would have its greatest effects around higher acceleration events, samples were created by looking for peaks of activity in each cat’s trace and selecting a fixed amount of activity data before and after the peak occurred, effectively building a window around the peaks (Fig. 1).

**Figure 1:**
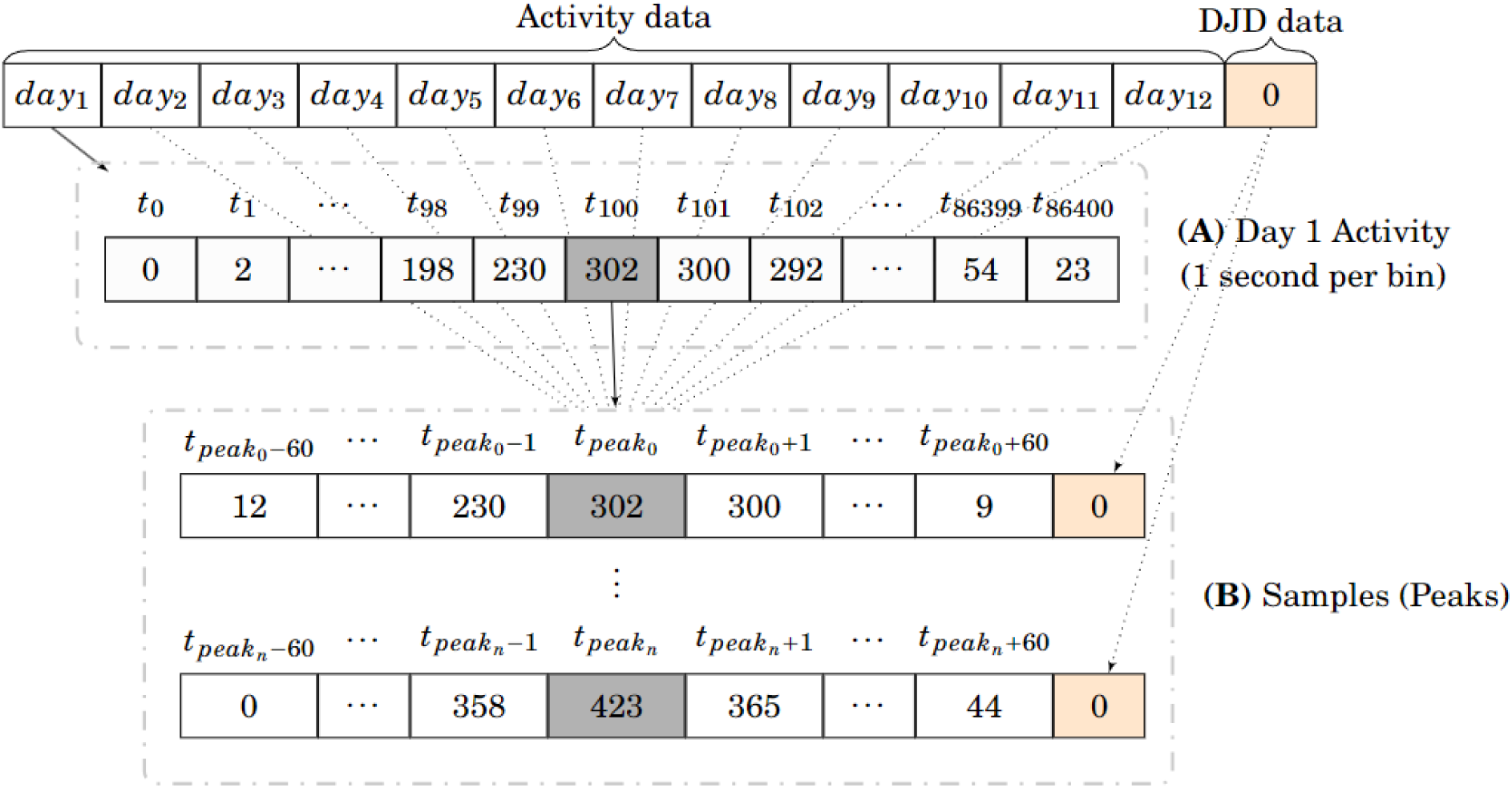
Schematic of the ML samples for a single cat. In this Illustration, we explain how we created samples for a single cat. For each cat, an activity count was recorded every second for 12 days and the cat was assigned a DJD health status (0 for healthy, 1 for unhealthy) by its owner. (A) shows the activity data of the first day from the first second to the last second of the day. Our approach looks for peaks of high activity, in this example, the first peak (peak0) was found in Da y1 at t = 100, we will create a sample by selecting the activity in the window centred on t100 = tpeak0 of length 2 minutes (120 seconds), the peak created will be assigned the health of the animal, in this case healthy. By following this process we can build an array of n samples containing the activity data around the top n peaks (B). Note that a single day of activity can contain multiples or no peaks, for this example the first day only contains a single peak.

This also had the effect of aligning the samples (i.e. by the point of impact, Fig. 2). The window length was fixed to different values (15, 30, 60 and 160 seconds) and the top n (Where n was variable) most intense peaks were selected based on the activity count.

**Figure 2:**
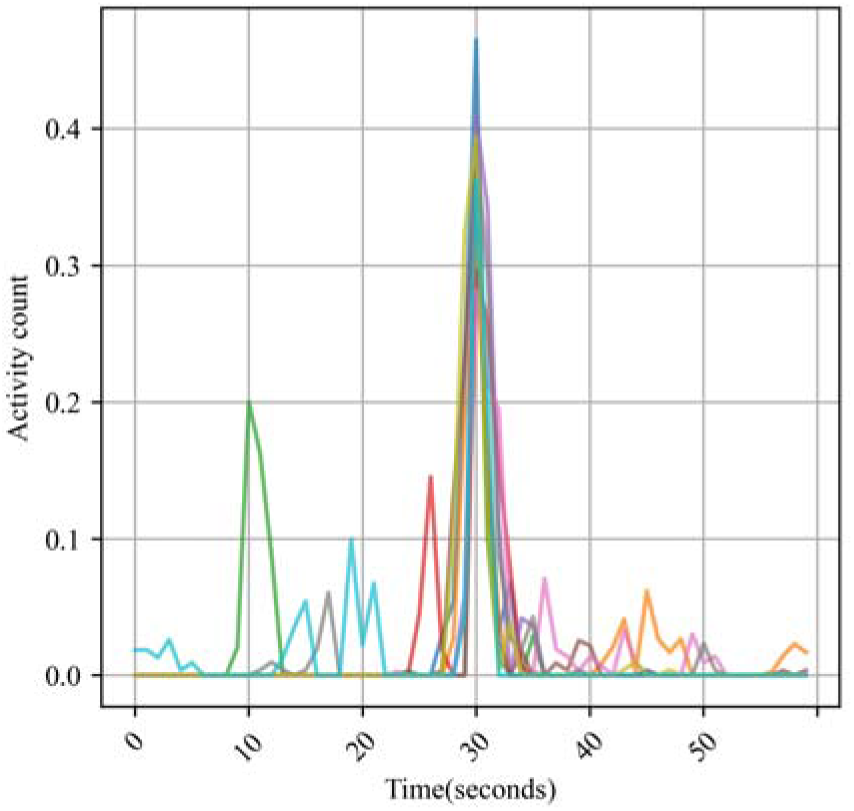
1-Peak samples for a single healthy cat. Shows 60 seconds 1-Peak samples for a single healthy cat. The x-axis shows the time in seconds (length of the sample) the y-axis shows the activity count value.

Data augmentation is a technique used in machine learning to increase the size of a dataset by creating new, synthetic data points that are similar to the original data points with the goal of increasing the diversity of the data available to the model. Small datasets are subject to overfitting (i.e. when a model adjusts excessively to the training data and is unable to predict unseen data correctly); one solution is data augmentation that increases the number of samples from existing data (Liu et al, 2020, Um et al 2017). Combinations of peaks from the initial sample dataset were used to augment the data in the present study (Fig. 3., Fig. S4). The number of possible combinations was capped at a hundred per cat to limit computational expense. We train our models with and without augmentation of the data. For the augmented datasets we use 2 to 25 peak combinations.

**Figure 3:**
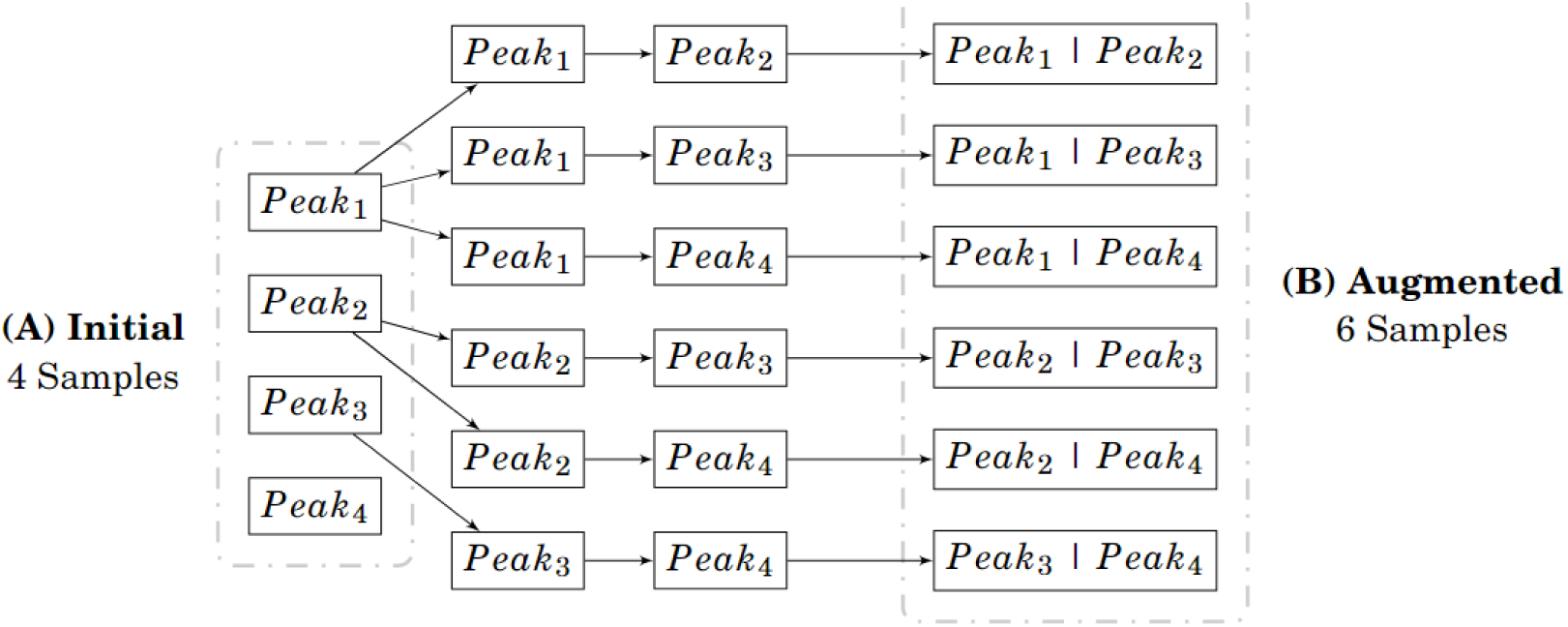
Cat samples augmentation with combinations for a single cat. By using per- mutations we can increase the number of samples from 4 samples containing 1 peak each (A) to 6 samples containing 2 peaks each (B).

**Figure 4:**
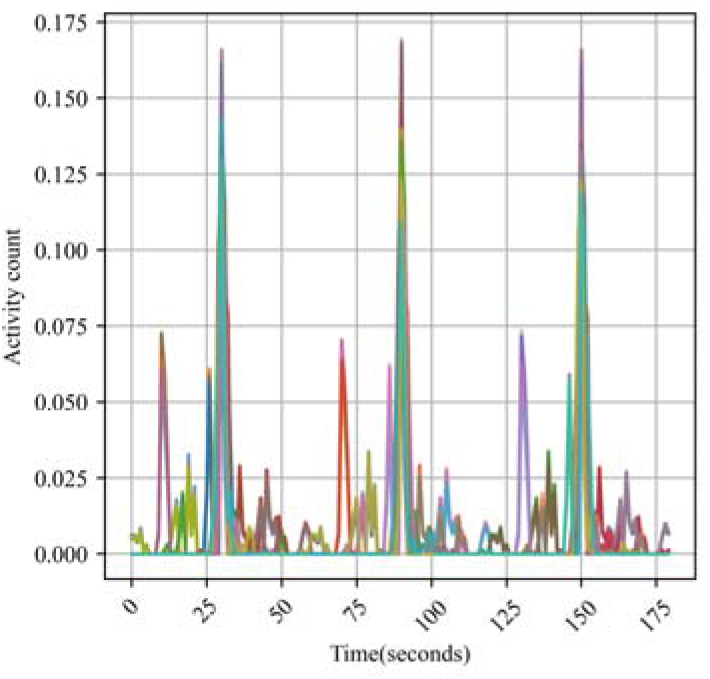
3-Peak samples for a single healthy cat. Shows the augmented version of (Fig. 2) by combinations of 3 peaks. The x-axis shows the time in seconds (length of the sample) the y-axis shows the activity count value.

### Pre-processing

Although the collar positioning was the same for all individual, cats have unique anatomical differences, which can cause variations in how a collar-mounted sensor is positioned.

Therefore, to compare activity patterns within each sample, the data were aligned and normalized to account for these slight positional differences. L1 normalisation (Bourbaki N, 1987) was used to scale individual data points within a dataset such that the sum of their absolute values equals one. This normalisation method helps emphasise the relative importance of features by ensuring that their contributions are weighted equally in terms of magnitude. The following pre-processing approach was used for the activity count data; firstly, the normalised sample was scaled with the original raw median sample to restore the natural range of the activity data and prepare for the Anscombe transformation followed by logarithmic transform, which transforms Poisson-distributed count representation of a log-normal activity signal to follow a normal distribution which most machine learning methods assume (Fig. 5). In addition, the log transform enables us to focus on proportionate rather than absolute changes in activity for each cat (Refer to the supplementary material for further details).

**Figure 5:**
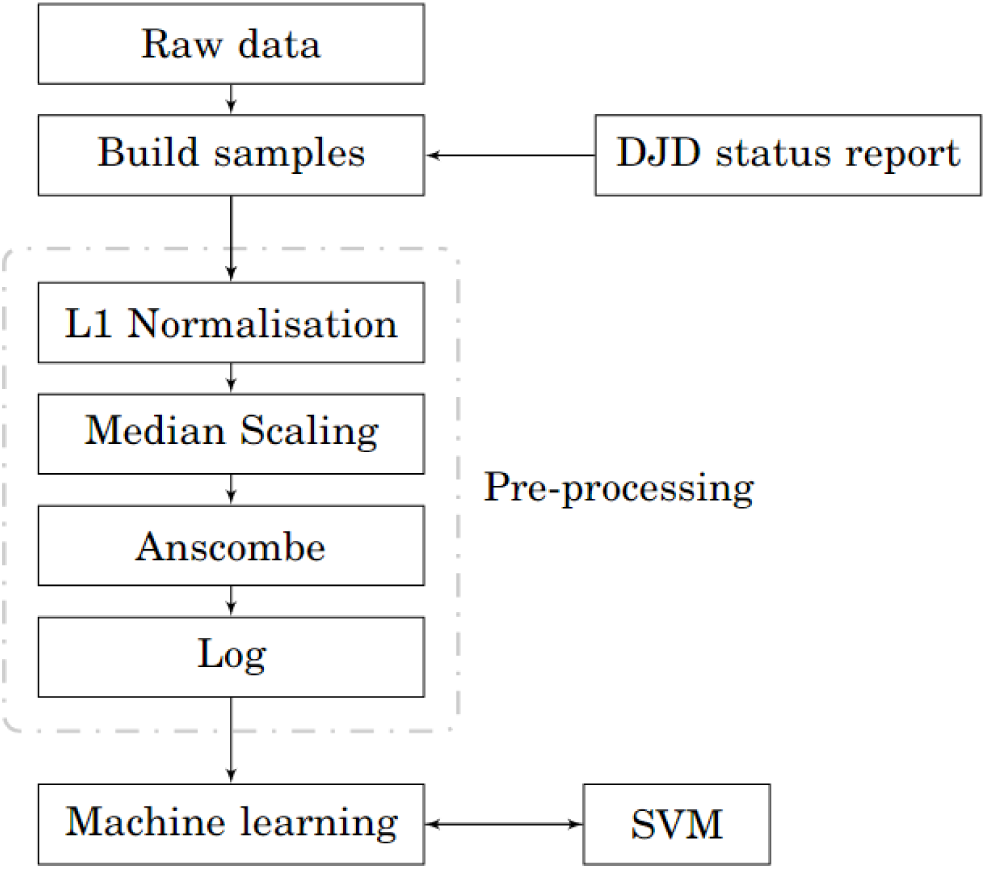
Classification of health status pipeline. Diagram showing the pipeline used for the classifying of healthy from unhealthy samples.

### Evaluation

#### Leave one out cross-validation

Data leakage is a common problem in ML where information from the test set is inadvertently used to train the model. Since our augmentation approach duplicates data across multiple samples from the same cat, it’s crucial that identical peaks do not appear in both the training and test sets, and indeed, that the same cats are not used for both training and testing. Unlike conventional leave-one-out cross-validation (LOOCV) (Wong, Tzu-Tsung, 2015), which is performed at the datapoint level, we adopted a cat-level LOOCV. This means that the model is trained on all cats except one, and then tested on the cat that was left out, ensuring that each cat is used for testing exactly once (Fig. S3). This process is repeated for each observation in the dataset, and the results are aggregated to give an estimate of the model’s performance.

### Area under the curve, receiver operator characteristic curve and precision

Receiver operator characteristic (ROC) curves and the area under the curve (AUC) were used to evaluate the performance of the classification models. These metrics rely on a confusion matrix, which summarises how well the model distinguishes between different classes, such as healthy and unhealthy animals. The ROC curve illustrates the relationship between the true positive rate (sensitivity) and false positive rate (1 - specificity) across various threshold values used for classification. A higher AUC value, ranging from 0 to 100%, indicates better discrimination ability of the model, whereas 50% means no better than random chance. By providing a visual representation and a single scalar metric, ROC curves and AUC facilitate the comparison of different models and help in determining the optimal threshold for classification tasks (Hoo et al, 2017) (refer to Supplementary material for additional details).

### Bootstrapping for confidence intervals

Due to the use of LOOCV, each test split only contains the samples from a healthy or unhealthy cat, which does not permit to create the confusion matrix necessary for the ROC curve. A single test ROC curve was therefore created by concatenating the predicted and true health status labels of all the cats used for testing. The process of bootstrapping involves randomly selecting a subset of the original dataset, with replacement, to create a new “bootstrap sample”. This means that each data point in the original dataset has an equal chance of being selected multiple times in the new sample. This bootstrap sample has (approximately) the same statistical properties as the original dataset and could have been generated instead of our original dataset. This process is repeated multiple times to create several new bootstrap samples. By passing each sample through the same analysis we can then determine the different the effect random chance has on our results. Bootstrapping can be performed at any point in the pipeline but applying it before machine learning would require the whole LOOCV machine learning to be performed 1000 times, once for each bootstrap sample. While this would have captured all uncertainty, it would have also biased the performance of the ML downwards by effectively reducing the sample size further for what is already a small study. In this study, bootstrapping was applied to the output of a single LOOCV ML analysis of the original dataset. One thousand bootstrap samples were therefore generated on the predictions of each unique fold obtained by LOOCV, which were then sorted to determine the 95% quantiles, which correspond to the 95% confidence interval of the model’s results (Efron et al, 1994).

### Sensitivity and Specificity

The optimal sensitivity and specificity were calculated by using the Youden index (Perkins et al). Due to the use of LOOCV, the sensitivity was calculated by using all the ground truth and all the predictions together.

### Support Vector Machine

In addition to SVM, we also explored other classification models, including K-nearest neighbours (KNN), Decision Tree (DTREE), and Logistic Regression (LREG), with results detailed in the supplementary material (Table S3 and Fig. S4). SVM was selected for this study as it demonstrated the best performance among these classifiers.

SVM (Boser et al) is a widely used method in machine learning for classifying data. It works by finding the best way to separate data points into their respective groups based on their characteristics.

When data points are not easily separable, SVM transforms the data into a higher-dimensional space where it is easier to find a dividing boundary. The goal is to find the boundary that maximises the margin, or distance, from the closest points of each category, ensuring a clear distinction between groups.

SVM can also handle cases where some classification errors are allowed, making it more flexible for real-world data. By using functions called kernels, such as the Radial Basis Function (RBF), SVM can create complex boundaries for better classification.

The performance of SVM depends on certain parameters, which are fine-tuned through cross-validation to achieve the best results. In essence, SVM is a powerful tool for accurately categorising data by finding optimal separating boundaries.

## Results

### Classification of DJD status with activity data

The best SVM model was trained on 30 second samples with combinations of 22 peaks and had a 78% median AUC with (0.65%, 0.88%) 95% confidence interval (Fig. 6). Each LOOCV fold was trained on 5000 samples and tested on 100 samples. The sensitivity was 68% (0.64%, 0.77%) and the specificity was 75% (0.68%, 0.79%).

**Figure 6:**
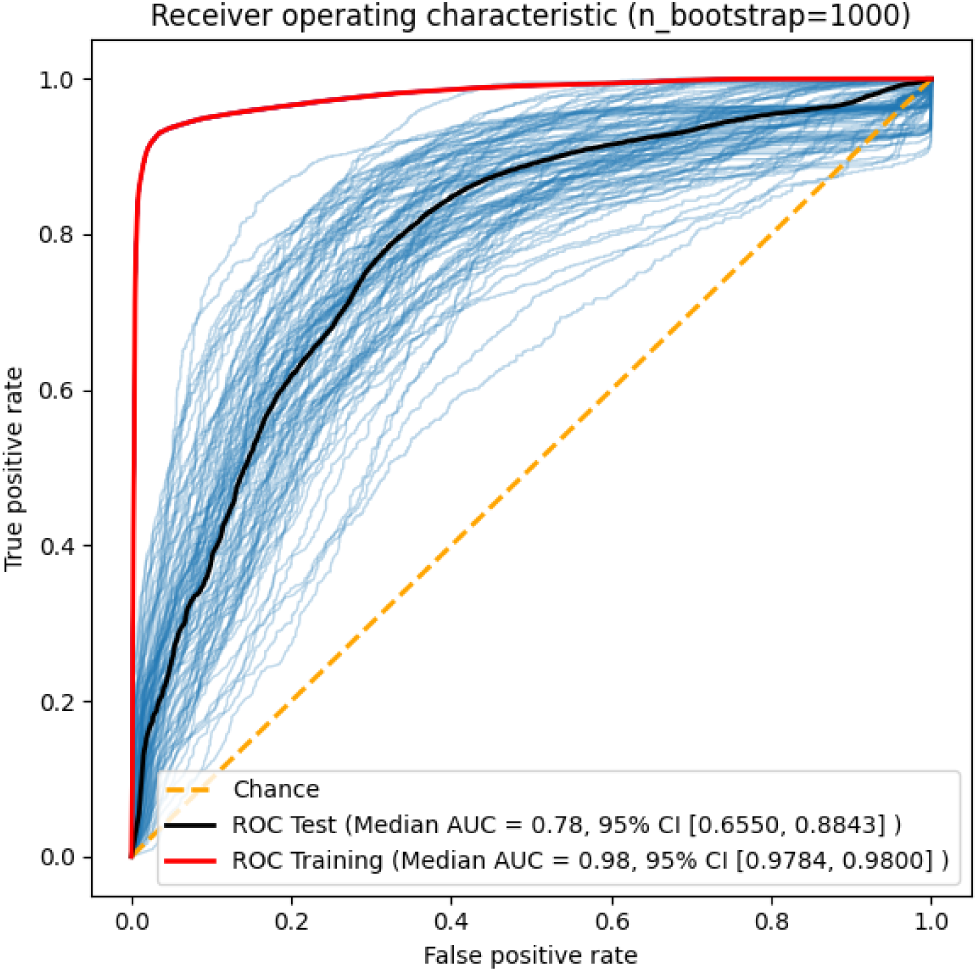
AUC and ROC curves of best SVM model. The red curve shows the training AUC while the black curve shows the testing AUC. The blue curves show the confidence interval.

### Augmentation and postprocessing of accelerometry data

To demonstrate the impact of our augmentation approach the sample length was fixed to 1 minute and no postprocessing was applied to the accelerometry data.

In Figure 7, the ROC curves are displayed for different datasets with and without data augmentation. In Figure 7A a 65% AUC was obtained with the 1-peak dataset (no data augmentation, 25 peaks per cat were generated, resulting in a total of 1275 samples. In Figure 7B a 20-peak combination was used which increased the AUC to 75%.

**Figure 7:**
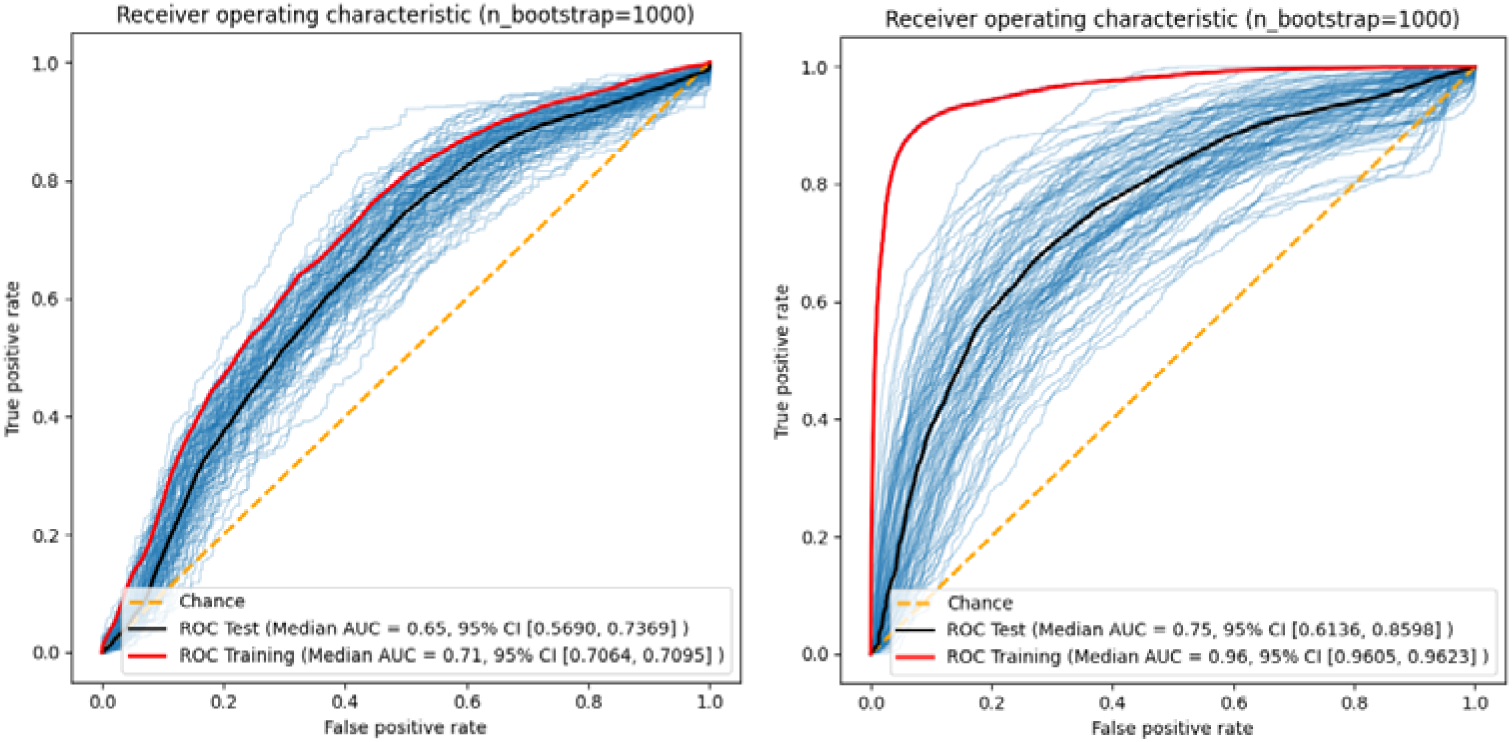
AUC and ROC curves when increasing the number of peaks. The red curve shows the training AUC while the black curve shows the testing AUC. The blue curves show the confidence interval. While (A) shows the non-augmented data ROC, (B) shows the augmented (22 Peaks).

Fig. 8 demonstrated the AUC performances against the augmentation amount with different sample lengths, 15, 30, 60 and 120 seconds for the SVM model. Each curve shows the training/testing AUCs for different accelerometry data pre-processing pipelines.

**Figure 8.**
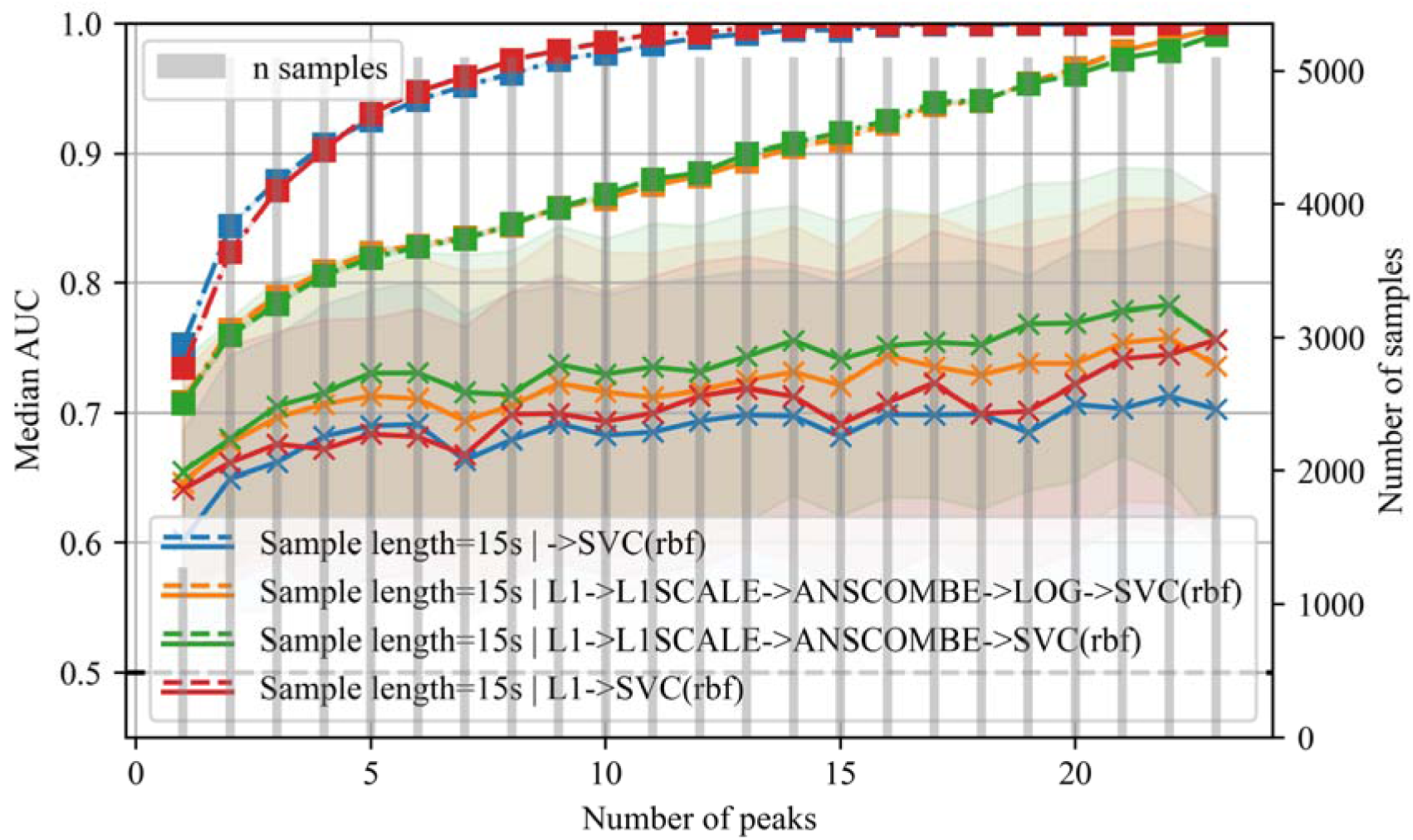
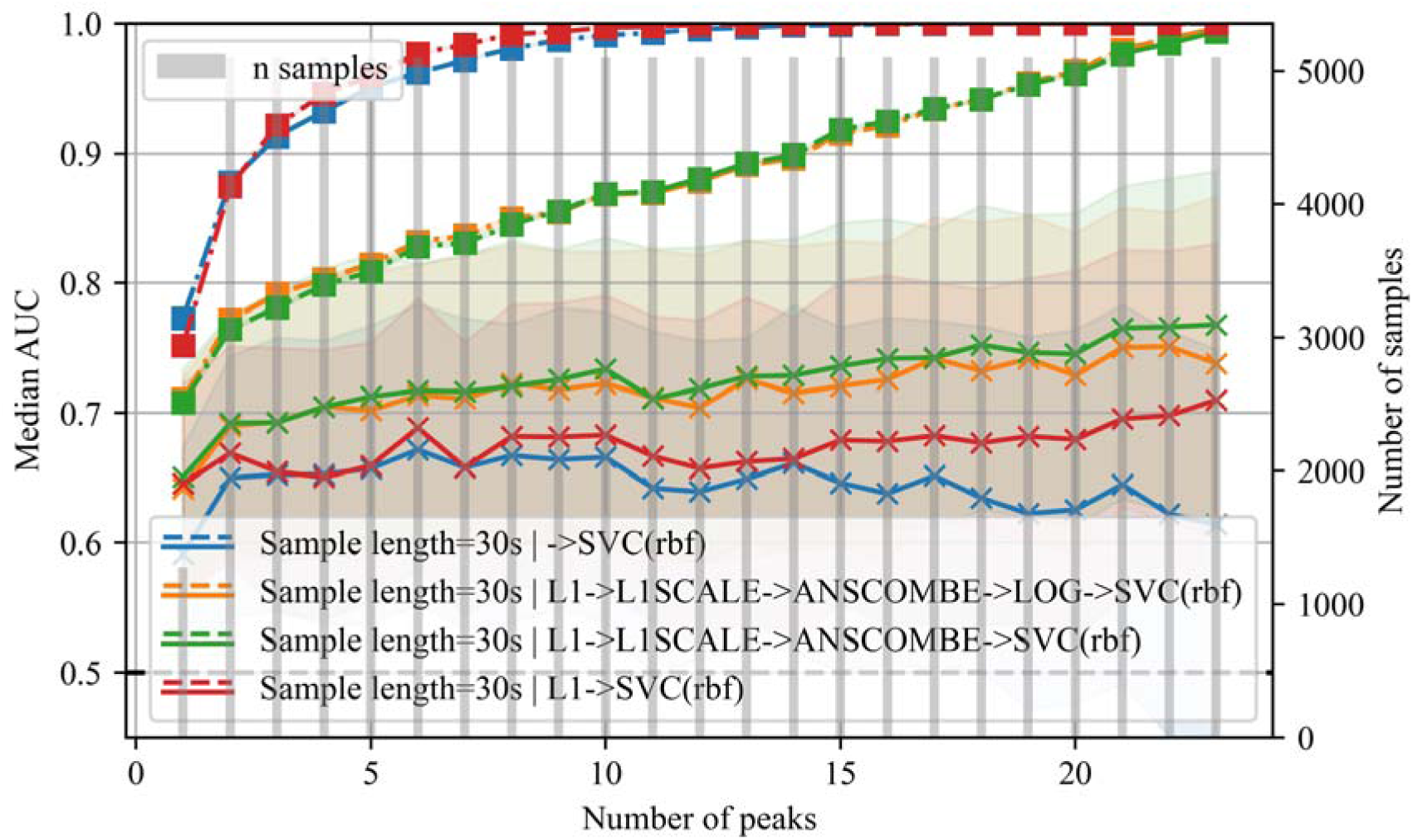
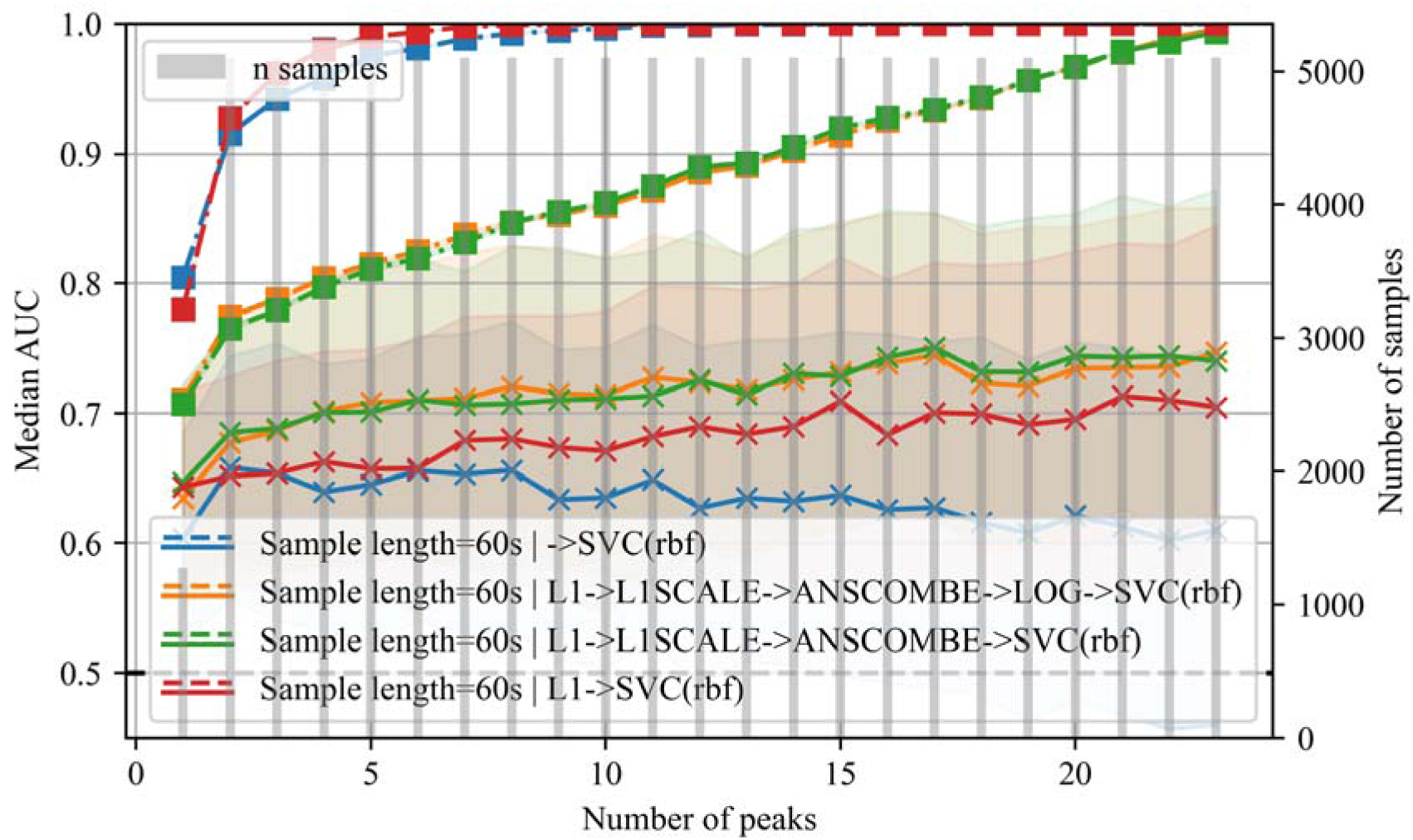
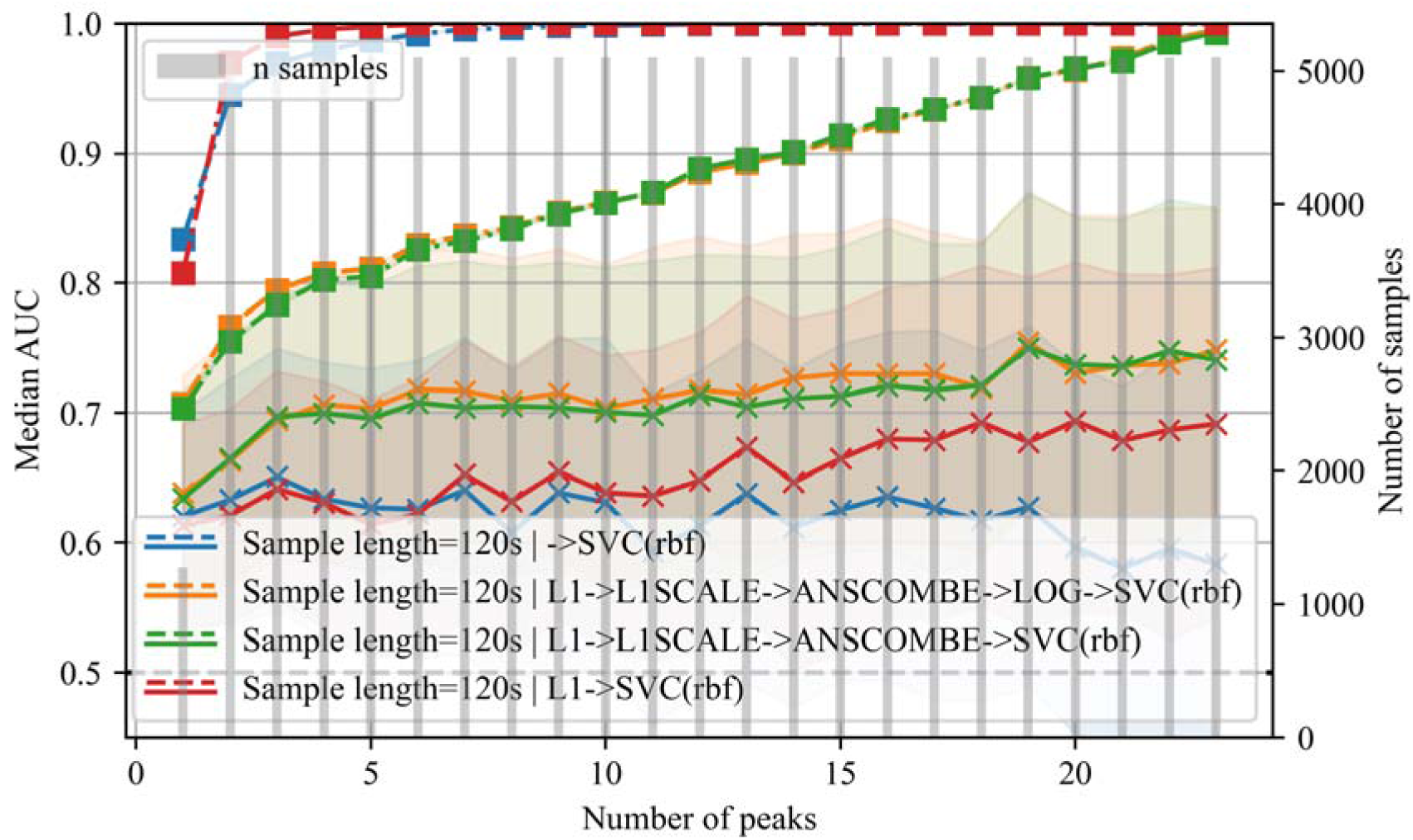
A, B, C, D, E: **Evolution of the AUC with the increase of the number of peaks**. While the x-axis shows the number of peaks used in the data set, the y-axis shows the Median AUC. The training AUC is shown by the dotted lines while the testing AUC is shown by the solid lines. Each subfigure is for the 15, 30, 60 and 120 seconds peak dataset respectively.

Augmenting the dataset increased the predictive power of the models, e.g. for a sample length of 30 seconds and when using L1 normalisation, Anscombe and logarithmic transformation (Fig. 8B), the non-augmented dataset had a 65% AUC that increased to 70% with the 2-peak dataset and 76% for 20-peak dataset; a similar increase was achieved with the 15, 60, and 120 second datasets (Fig. 8A, Fig 8C, Fig 8D).

Using L1 normalisation improved the model performances, e.g., Fig. 8B shows the AUC values of the 30-second datasets where we trained the model with the raw accelerometry data (without normalisation and no pre-processing) as a blue curve, while the red curve shows the results when using L1 normalisation. In addition in Fig. 9 we show the AUC difference between the normalised (red curve) and the non-normalised (blue curve) data, highlighting the improvement.

**Figure 9:**
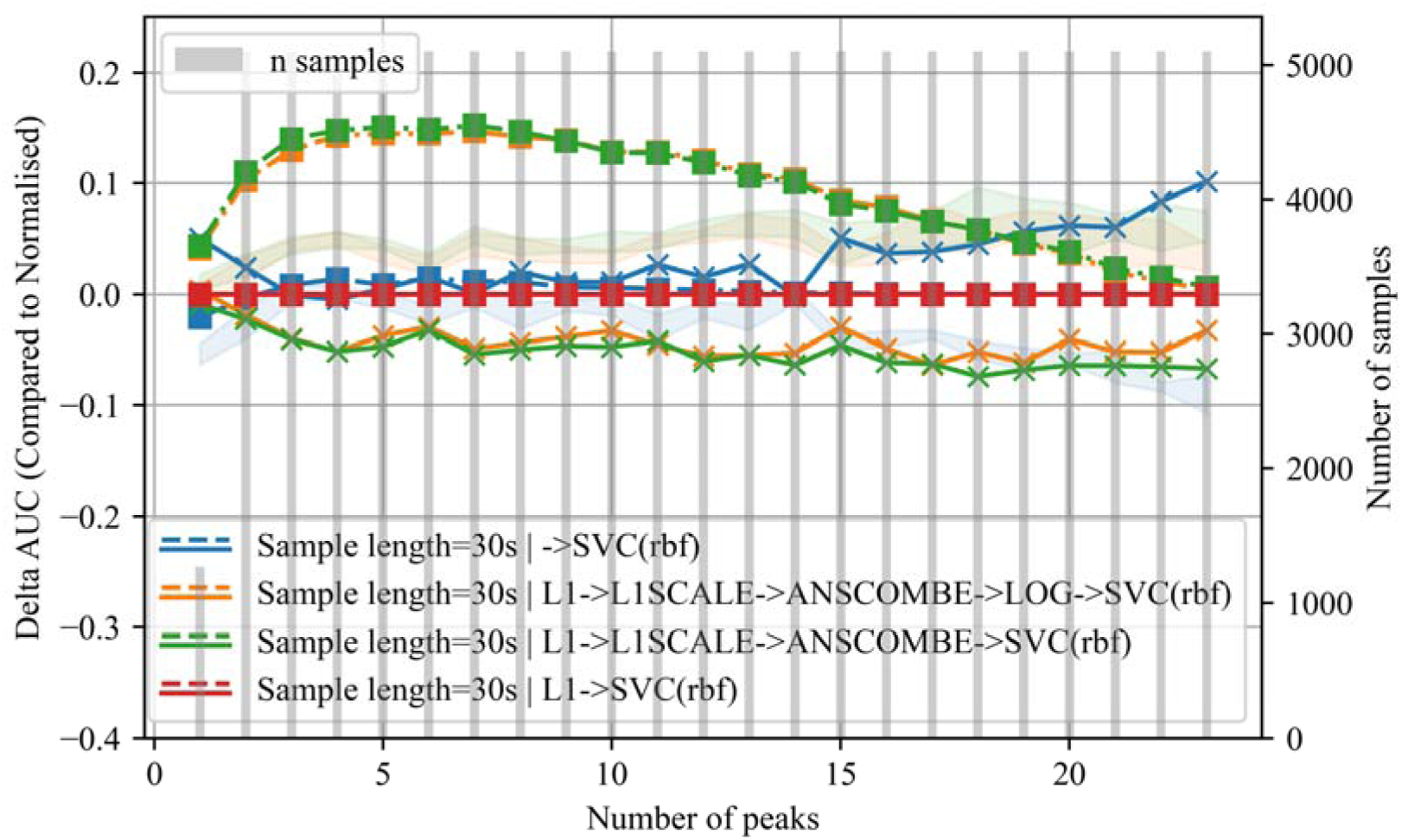
Impact of Normalisation on predictive power. While the x-axis shows the number of peaks used in the data set, the y-axis shows the difference between the AUC of the normalised datasets against the non-normalised.

The sample length does not significantly change the predictive power of the models, for all four sample lengths the 20-peak dataset with L1 normalisation, Anscombe and logarithmic transformation had an AUC of approximately 75% (Fig. 8).

### Classification of DJD status with age

DJD is known to be prevalent in older cats (Bennett et al, 2012). Models were therefore trained and validated using age only, to assess the predictive power of our accelerometry model. The AUCs were compared between these models using the Wilcoxon signed-rank test (Fig. 10).

**Figure 10:**
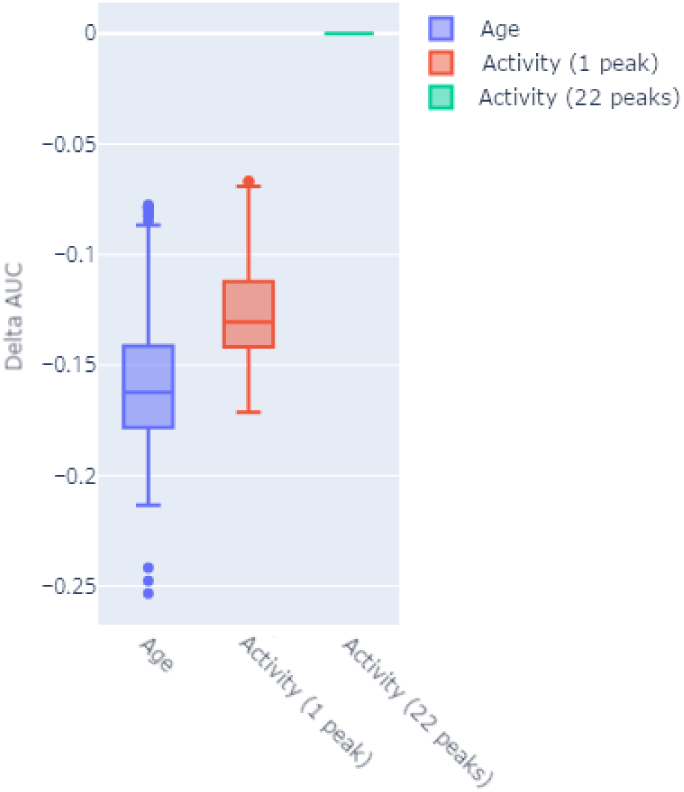
Performance of SVM model trained on activity against age. The y-axis shows the difference compared to the best model AUC (Activity with 22 peaks).

The AUC for the accelerometry data was significantly greater than that using age alone and demonstrated improved performance, although using a single peak of activity per cat is statistically significant, using multiple peaks increased performance further (Wilcoxon p<0.001),.

## Discussion

### Model explainability

After fitting the ML model, it is possible to visualise what parts of the data in the samples are most important for the classification process (Fig 11). Because the model was fitted with accelerometry time series it is possible to directly see what part of the time series is most useful for the model to discriminate between DJD and no DJD cats. In Figure 11A a sharp increase of feature importance could be observed approximately 7 seconds before the high activity event until 7 seconds after, which was also observable in the other augmented versions of the dataset (Fig. 11B, 11C). This pattern may be due to DJD cats preparing differently for the impact, and noticeably the most important features were just after impact, which may be due to cats recovering from the shock.

**Figure 11.**
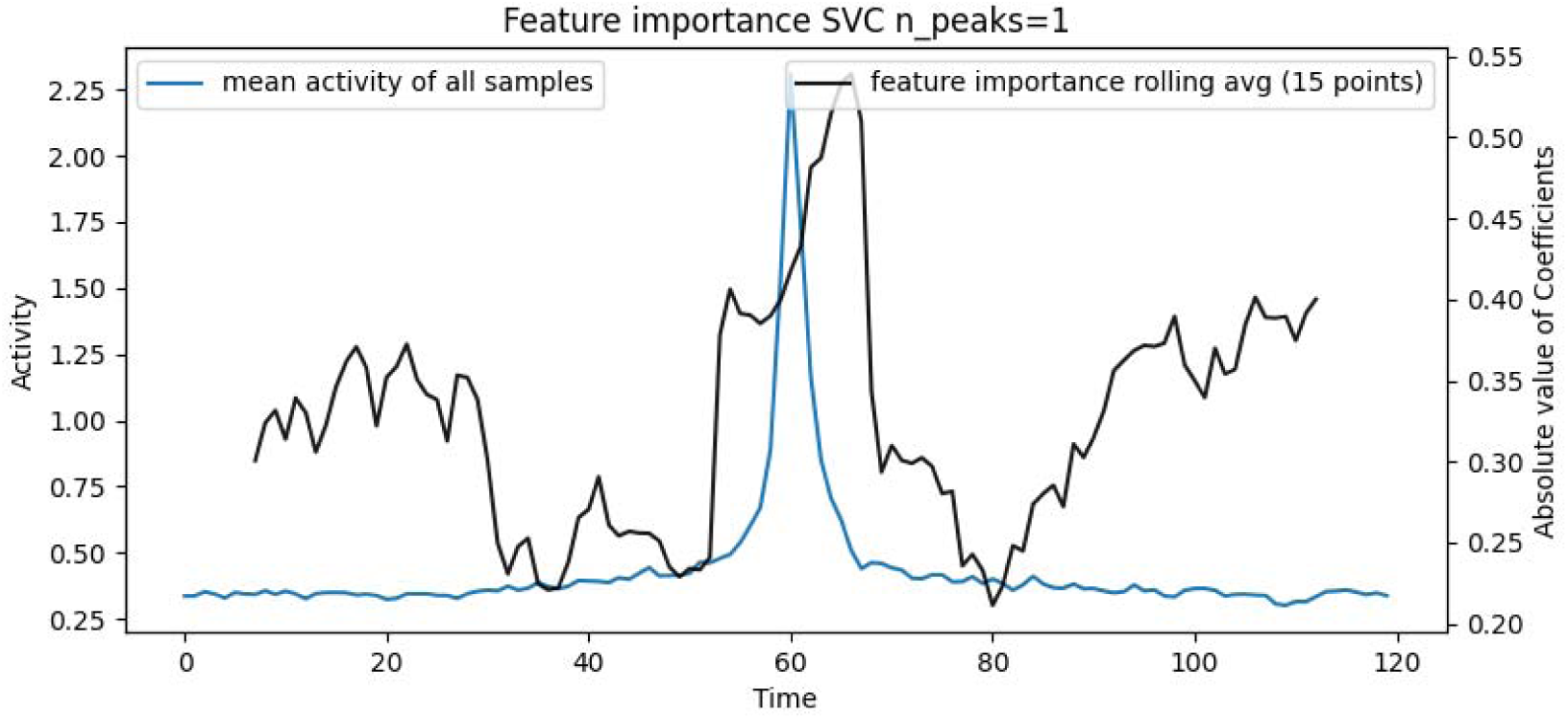
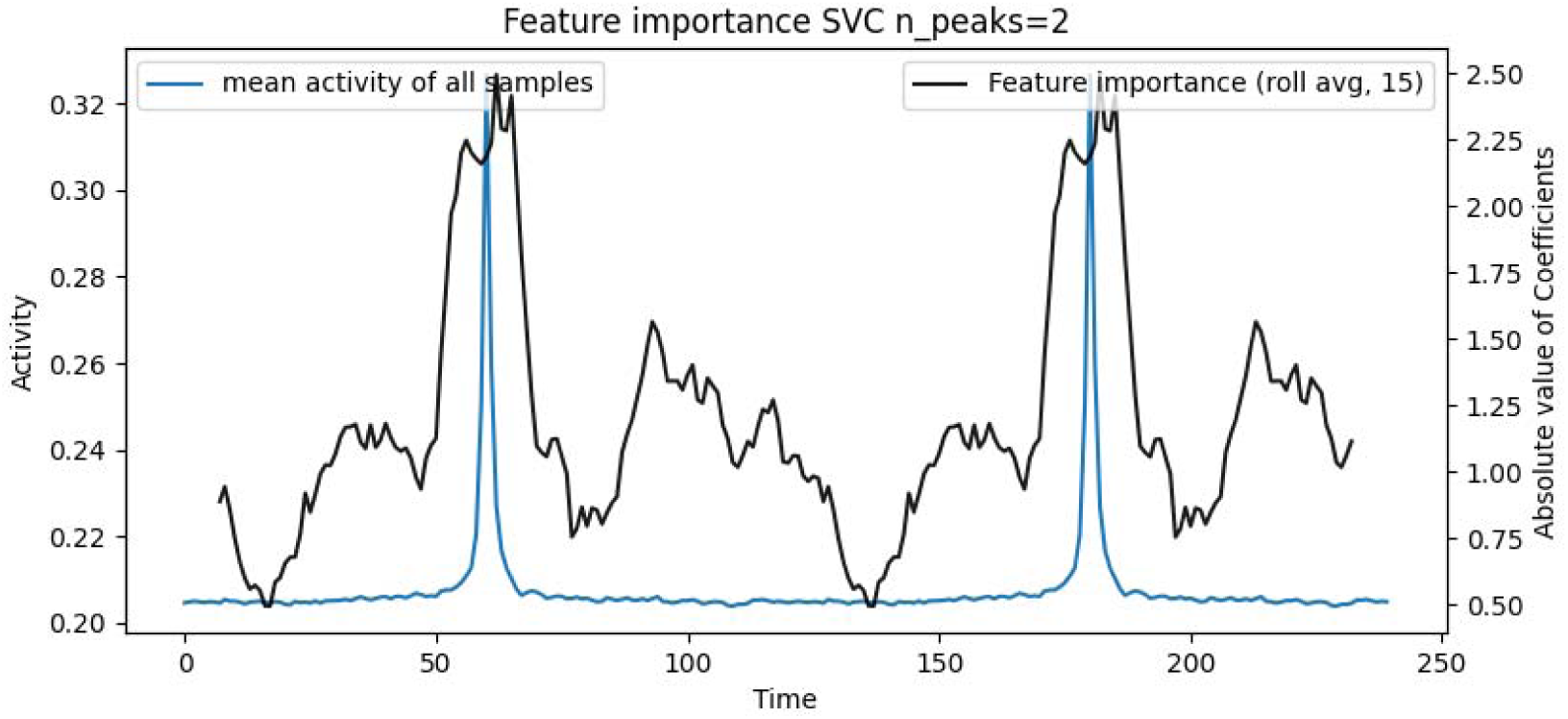
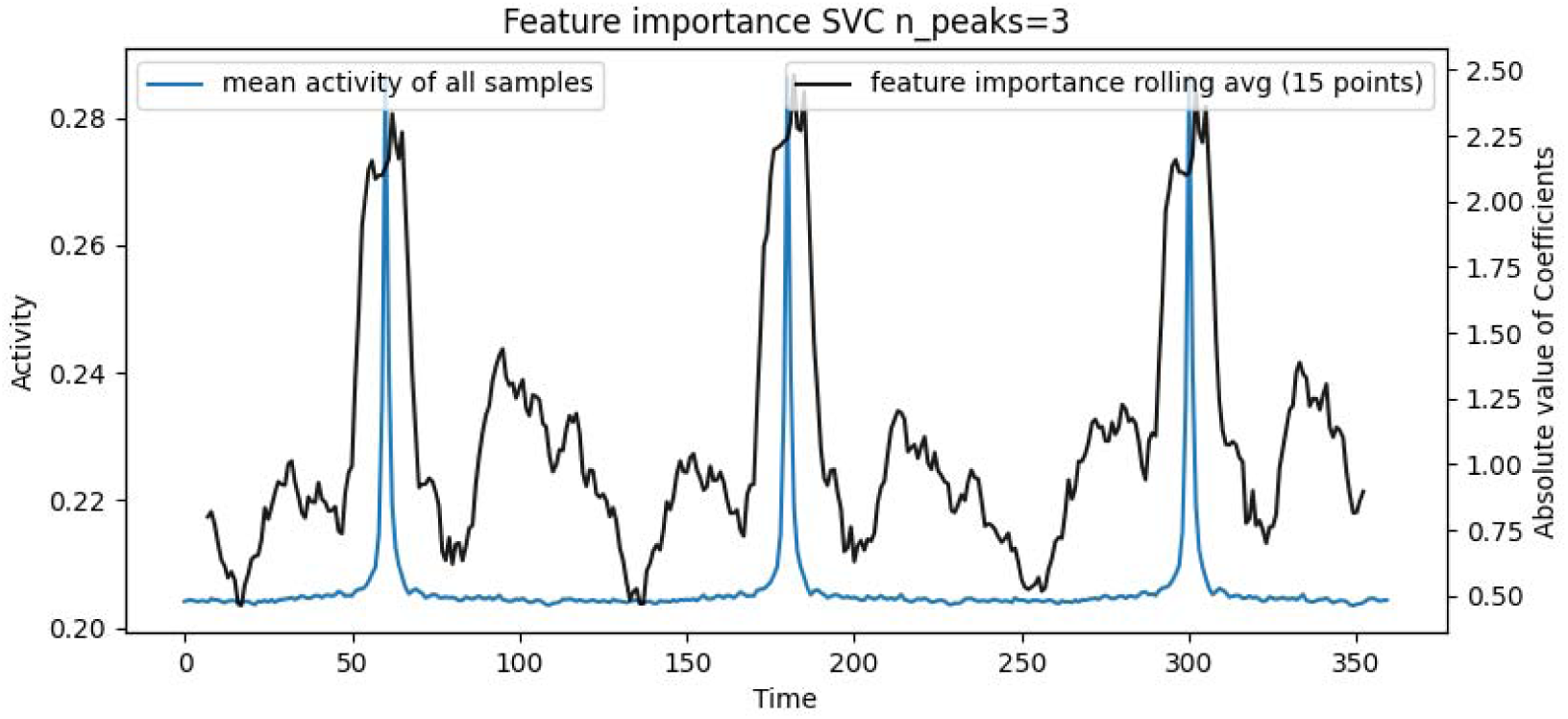
A, B, C: **Feature importance**. The Blue curve shows the mean trace of all the samples used to fit a linear SVM model. The Black curve displays the feature importance derived from the fitted model. While the x-axis shows the time, the y-axis on the left shows the mean activity count of all cats after normalisation and pre-processing, the y-axis on the right shows the SVM feature importance.

For this study, interpretability is key for veterinarians as it would allow them to devise appropriate solutions and treatments to alleviate negative clinical signs earlier and increase the well-being of the cats. Accelerometers mounted on collars were deployed on cats over 14 days to collect crucial activity data for predictive modelling, focusing on early DJD detection. The supervised machine learning approach, with SVM performing best, analysed temporal segments of heightened activity to correlate patterns with DJD indications. By constructing samples around high-activity peaks, DJD effects during intense movements were captured.

Model evaluation through LOOCV demonstrated promising results, with SVM models achieving a median AUC and sensitivity of 78% in classifying DJD status. The study underscores the efficacy of accelerometer data over age in predicting DJD, emphasising the potential clinical utility of activity monitoring. In clinical settings this could involve user-friendly tools that simplify data analysis and interpretation. For example, automated systems could provide straightforward visual representations via monitoring software, aiding clinicians in making informed decisions about patient care. Such tools could streamline the process of assessing cat joint health, potentially leading to earlier detection and intervention for DJD.

Our findings align with previous studies that emphasise the significance of monitoring physical activity for detecting DJD (Gruen et al, 2017; Yamazaki et al, 2020). However, this study goes further by providing a robust statistical analysis and machine learning validation of accelerometer data. The use of LOOCV at the cat level ensured that the model’s performance was evaluated rigorously, avoiding data leakage and providing a more accurate assessment of its predictive capabilities.

Interestingly, the study revealed that activity just before and after high-activity peaks is crucial for classification. This finding may indicate that cat apprehension influences their movement patterns before engaging in intense activities, possibly due to anticipated pain. This insight suggests that not only the intensity of the activity but also the behavioural context of the activity is important for identifying DJD.

This study has a number of limitations that should be considered. First, the sample size of 51 cats, although substantial, may not fully capture the variability seen in the general cat population. Future studies with larger, more diverse populations would provide a more robust validation of the findings. Additionally, the study duration of 14 days, while sufficient to capture activity patterns, might not account for longer-term fluctuations in activity levels due to external factors such as changes in the home environment or the cat’s health status over time. Expanding the monitoring period could offer deeper insights into the chronic progression of DJD.

Moreover, although collar positioning was the same the use of accelerometers mounted on collars may introduce variability in data collection due to slight differences in collar fit.

Ensuring consistent placement and securing of the devices is crucial for accurate data acquisition. Another consideration is the reliance on owner-reported mobility evaluations, which can be subjective and may vary in accuracy. Incorporating more objective assessment tools, such as video analysis could complement owner reports and enhance the reliability of mobility assessments.

While the SVM model showed promising results, it is essential to explore and compare other machine learning algorithms to determine the most effective approach for DJD detection, we compared SVM against KNN, DTREE and LREG (Supplementary material) but other classifiers could yield better results. Particularly deep learning models can handle intricate data relationships through multiple layers of abstraction, potentially offering superior performance for DJD detection but will require substantially greater amounts of input data for training.

By relying solely on the wearable accelerometer, researchers can avoid the complexities of collecting external data such as weight, breed, and medical history, simplifying the study while still achieving valuable insights directly from the device itself. However future research should also investigate the integration of such additional features.

## Conclusions

The machine-learning approach described in the present study, supported by data augmentation, shows promise in predicting early joint disease in cats. The analysis focused on small windows around activity peaks revealing discriminative information distinguishing cats with and without owner-reported signs of DJD. Ongoing research aims to identify critical activity segments for prediction, offering insights into how cats with early DJD differ in their movements during high-intensity actions. While this method uncovered discriminative markers in activity zones, it is not entirely generalisable across a cat’s entire activity trace.

Building a health classifier will necessitate incorporating these findings within a broader model. Data augmentation aids instead of large datasets yet expanding participant numbers remains pivotal for robust machine learning models. Early findings encourage further exploration or expansion of this study for more comprehensive insights.

## Supporting information

Questionnaire Form

## Acknowledgments

We are grateful to the cat owners for their participation and to the University of Bristol’s ethics committees for approving this study. Their collective contributions were essential for advancing the early detection of DJD in cats.

## References

Boyd A, Golding J, Macleod J, Lawlor DA, Fraser A, Henderson J, Molloy L, Ness A, Ring S, Davey Smith G, 2013. Cohort Profile: the ’children of the 90s’--the index offspring of the Avon Longitudinal Study of Parents and Children. Int J Epidemiol. 42(1):111–27.

Anscombe, F.J, 1948. The transformation of Poisson, binomial and negative-binomial data. Biometrika 35, 246–254.

Benito J, Hansen B, Depuy V, Davidson G, Thomson A, Simpson W, Roe S, Hardie E, Lascelles B, 2013. Feline musculoskeletal pain index: responsiveness and testing of criterion validity. Journal of Veterinary Internal Medicine 27, 474–482.

Bennett D, Zainal Ariffin, S.M. bt, Johnston, P., 2012. Osteoarthritis in the cat: 1. How common is it and how easy to recognise? Journal of Feline Medicine and Surgery 14(1), 65–75.

Boser B.E, Guyon I.M, Vapnik V.N, 1992. A training algorithm for optimal margin classifiers, 144–152.

Bourbaki N, 1987. Topological Vector Spaces: Chapters 1–5. Éléments de mathématique.

Davies V, Scott E.M, Wiseman-orr M.L, Wright A.K, Reid J, 2020. Development of an early warning system for owners using a validated health-related quality of life (HRQL) instrument for companion animals and its use in a large cohort of dogs. Frontiers in Veterinary Science 7, 575795.

Efron, Bradley, and Robert J. Tibshirani. An introduction to the bootstrap. Chapman and Hall/CRC, 1994.

Griffin FC, Mandese WW, Reynolds PS, Deriberprey AS, Blew AC, 2021. Evaluation of clinical examination location on stress in cats: a randomized crossover trial. Journal of Feline Medicine and Surgery. 23(4), 364–369.

Gruen M.E, Alfaro-Córdoba M., Thomson A.E., Worth A.C, Staicu A.-M., Lascelles B.D.X., 2017. The use of functional data analysis to evaluate activity in a spontaneous model of degenerative joint disease associated pain in cats. PloS One 12.

Hoo Z.H., Candlish J. and Teare D, 2017. What is an ROC curve? Emergency Medicine Journal. 34(6), 357–359.

Katti S.K., Rao A.V, 1968. Handbook of the Poisson distribution. Technometrics 10, 412– 412.

Lascelles B.D.X, 2010. Feline degenerative joint disease. Veterinary Surgery 39, 2–13.

Lascelles B.D.X, Dong Y.H, Marcellin-Little D.J, et al, 2012. Relationship of orthopedic examination, goniometric measurements, and radiographic signs of degenerative joint disease in cats. BMC Veterinary Research 8, 10.

Lascelles B.D.X., Depuy V, Thomson A, Hansen B, Marcellin-Little D, Biourge V, Bauer J, 2010. Evaluation of a therapeutic diet for feline degenerative joint disease. Journal of Veterinary Internal Medicine 24, 487–495.

Liu B, Zhang Z, Cui R., 2020. Efficient time series augmentation methods, in: 2020 13th International Congress on Image and Signal Processing, Biomedical Engineering and Informatics (CISP-BMEI), 1004–1009.

Maharana K., Mondal S., Nemade B., 2022. A review: data pre-processing and data augmentation techniques. Global Transitions Proceedings.

Maniaki E, 2020. Risk factors, activity monitoring and quality of life assessment in cats with early degenerative joint disease.

Maniaki E, Murrell J, Langley-Hobbs S.J, Blackwell E.J, 2021. Associations between early neutering, obesity, outdoor access, trauma and feline degenerative joint disease. Journal of Feline Medicine and Surgery 23(10), 965–975.

Maniaki E, Murrell J, Langley-Hobbs S.J, Blackwell E.J, 2023. Do owner-reported changes in mobility reflect measures of activity, pain and degenerative joint disease in cats? Journal of Feline Medicine and Surgery 25(6).

Perkins N.J, Schisterman, E.F, 2005. The Youden Index and the optimal cut-point corrected for measurement error. Biom. J. 47, 428–441.

Slingerland L, Hazewinkel H, Meij B, Picavet P, Voorhout G, 2011. Cross-sectional study of the prevalence and clinical features of osteoarthritis in 100 cats. The Veterinary Journal 187, 304–309.

Um T.T, Pfister F.M, Pichler D, Endo S, Lang M, Hirche S, Fietzek U, Kulić D, 2017. Data augmentation of wearable sensor data for Parkinson’s disease monitoring using convolutional neural networks, Proceedings of the 19th ACM International Conference on Multimodal Interaction, 216–220.

Wong T.-T, 2015. Performance evaluation of classification algorithms by k-fold and leave-one-out cross validation. Pattern Recognition 48(9), 2839–2846.

Yamazaki A, Edamura K, Tanegashima K, Tomo Y, Yamamoto M, et al. 2020. Utility of a novel activity monitor assessing physical activities and sleep quality in cats. PLOS ONE 15(7): e0236795.

