## Supplementary Material for "Accelerometer-derived classifiers for early detection of degenerative joint disease in cats"

**Early disease detection classifiers for wearables on domestic cats**

**Study group**

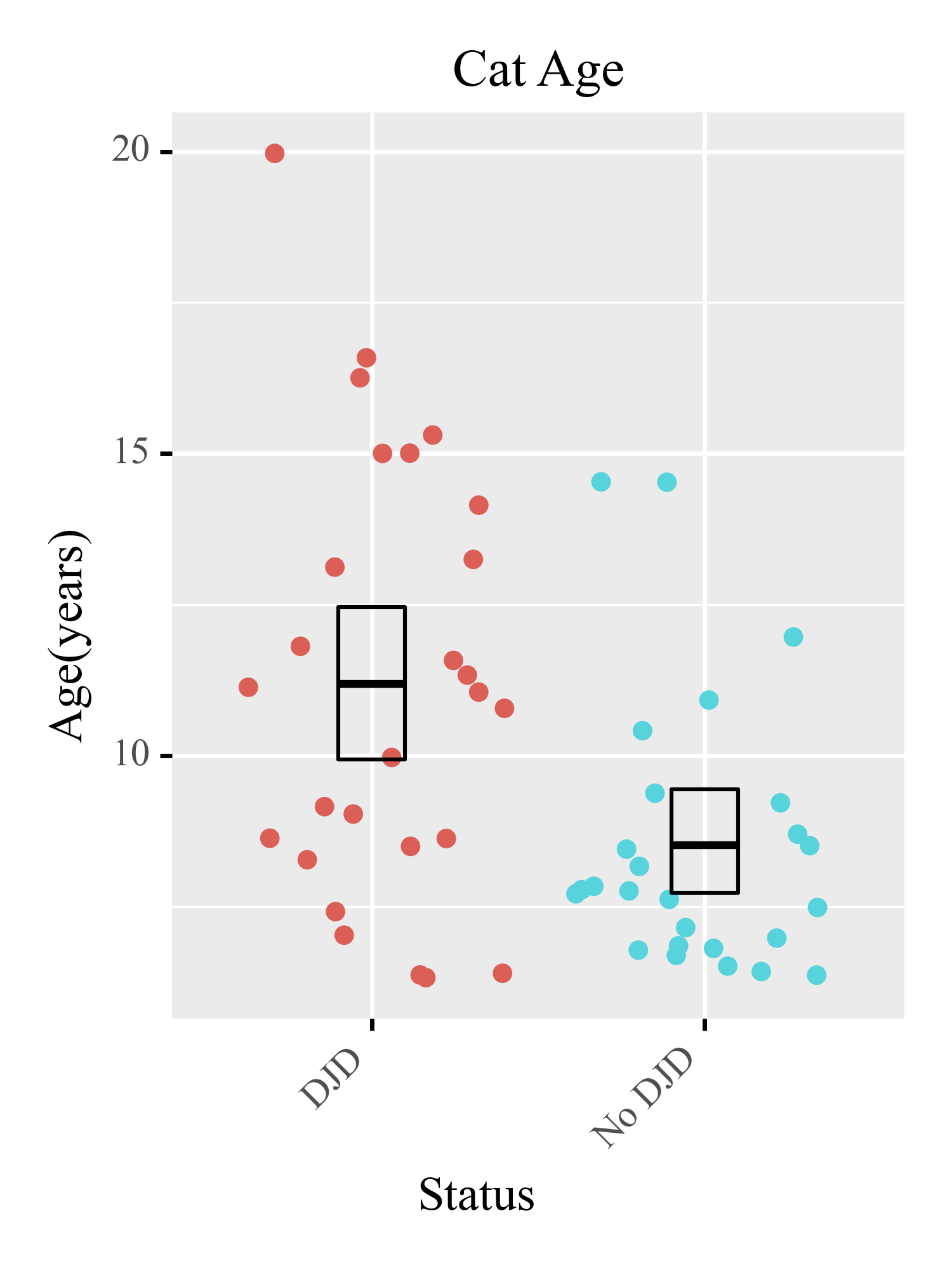

Figure S1.

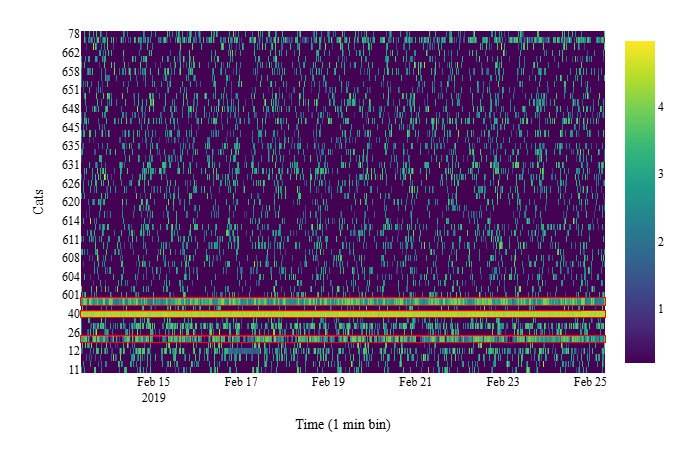

Figure S2.

**Methods**

*Anscombe*

The Anscombe transform (Anscombe et al, 1948) is a statistical data variance stabilisation transformation, i.e. a transformation which aims to change the input data so that the variance of each data point or observation is not related to its magnitude. This transformation therefore transforms heteroscedastic Poisson (Katti et al, 1968) distributed data into approximately Gaussian distributed data with a homoscedastic standard deviation of one. This then allows us to appropriately apply ML algorithms that assume Gaussian variance, which is the vast majority of them. Because our activity data are counts and more closely follow Poisson statistics than the Gaussian distribution expected by conventional machine learning methodology we will apply the Anscombe transform in our pre-processing pipeline.

$$Anscombe\left( x \right)=2\times\surd\left( x+\frac{3}{8} \right)$$

*Log*

We want to detect relevant bursts of activity as they may contain key information. In our use case, the percentage change in activity is more important than the absolute change, i.e. an increase in activity from 10 to 20 counts (100% increase) is far more important than from 100 to 120 counts (a 20% increased) despite both being an increase of 20 counts. The Logarithm function is a mathematical device that turns multiplicative change into additive change so that the ML can consider multiplicative effects symmetrically. Note that applying the Log transform on the output of the Anscombe transform does not cause problems because it is akin to a simple division by two except for a small attenuation at low counts caused by the 3/8 constant:

$$Log\left( x^{\frac{1}{2}} \right)=\frac{\log\left( x \right)}{2}$$

Five sensors were set up to measure activity counts every millisecond while the rest collected activity counts every second. For this project, all the activity data were resampled to the second resolution (i.e. each count contains the sum of activity counts within each second) if it was not already at that resolution. In Figure 3 we show the activity data (after Anscombe and Log transformation) of all the cats including the ones not used in the analysis. Each row shows the activity count data of a single cat during the entirety of the study.

**Area under the curve, Receiver operator characteristic curve and Precision**

To explain Receiver Operator Characteristic curves (ROC) and the area under the curve (AUC)

we must first introduce the concept of a confusion matrix. The confusion matrix summarises

how a model performs on the test data, the rows in a confusion matrix correspond to what the

ML algorithm predicted (for example, healthy and unhealthy) and the columns correspond to

the known truth (Table. S2) The true positives (TP) are defined as samples that were unhealthy

that were correctly identified by the prediction, the true negatives (TN) are the samples that

were healthy that were correctly classified, while the false negatives (FN) are when the sample is unhealthy but the model predicted it was healthy, and false positives (FP) are samples that were healthy but the model predicted them as unhealthy. ML algorithms output scores between -infinity and infinity of a given sample to be healthy or unhealthy, although many are able to convert this to a probability between 0 and 1. To make a decision a threshold must be set, for example, if we pick a 0.5 threshold all samples returning a probability above 0.5 will be categorised as unhealthy while those below 0.5 will be classified as healthy. By changing the value of the threshold for deciding if a sample is unhealthy or not

we change the calibration of the classifier and hence the values in the confusion matrix. For

example, if we want to prioritise the detection of unhealthy samples we can lower the threshold to 0.1 which will increase the number of TPs but also increase the number of FPs and also reduce the number of FNs. Similarly, we could set the threshold to 0.9 would decrease the number of FPs and increase the detection of TNs. By changing the threshold value we can find the optimal threshold value for our model for a particular task, ROC curves provide a simple way to summarise the effect of different threshold values. The y-axis shows the True Positive Rate (TPR), also called the sensitivity. The x-axis shows the False Positive Rate (FPR). The False positive rate tells us the proportion of healthy samples that were incorrectly classified.

We can calculate the TPR and the FPR for all of the confusion matrices generated by changing the value of the threshold from 0 to 1 and place the points on a ROC curve. A point at the top right corner (TPR = 1 and FPR = 1) means that even though we correctly classified all of the unhealthy samples, we incorrectly classified all of the healthy samples, while a point at the top left corner means that we correctly classified all unhealthy samples without mistake, the model makes "perfect" classifications. The line that goes through the points (0,0) and (1,1) is the "Chance line" where TPR = FPR any points on this line means that the proportion

of correctly classified unhealthy samples is the same as the proportion of incorrectly classified

healthy samples. The ROC curve summarises all of the confusion matrices that each threshold

produced.

The Area Under the Curve (AUC) is a value between 0 and 100 (in percent) is a metric that

allow a simple comparison of different model performances, with higher AUC indicating better performances.

**Evaluation**

**Leave one out cross-validation**

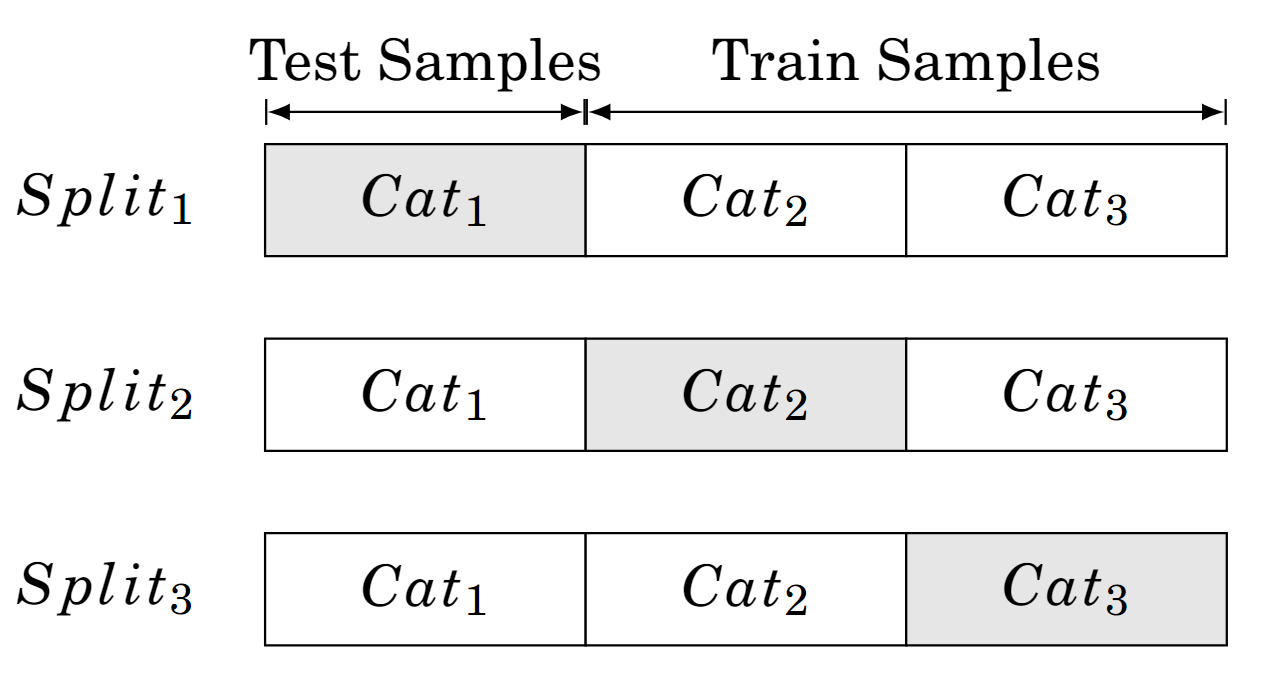

Figure S3.

**Regularisation**

We will conduct a grid search to identify the optimal parameters for our SVM model, exploring various combinations of kernel types, gamma values, and regularisation parameter (C) settings. This exhaustive search across the specified parameter grid aims to find the configuration that maximises the model's performance.

Similarly, for K-Nearest Neighbors (KNN), we will optimise by tuning the number of neighbors (k) and incorporating distance weighting to reduce overfitting, for Decision Trees (DTree), we will focus on regularisation by setting constraints on maximum depth, minimum samples per leaf, and for Logistic Regression (LReg), we will apply L1 regularisation.

**Bootstrapping for confidence intervals**

Receiver operating characteristic (ROC) curves are powerful tools for evaluating ML performance (See Appendix “Area under the curve, Receiver operator characteristic curve and Precision “). Because we used LOOCV, each test split only contains the samples of a healthy or unhealthy cat, which does not permit to create the confusion matrix necessary for the ROC curve. We therefore opted to create a single test ROC curve by concatenating the predictions and true health status labels (ground truth) of all the cats used for testing. The process of bootstrapping involves randomly selecting a subset of the original dataset, with replacement, to create a new "bootstrap sample". This means that each data point in the original dataset has an equal chance of being selected multiple times in the new sample. This bootstrap sample has (approximately) the same statistical properties as the original dataset, and could feasible have been generated instead of our original dataset. This process is repeated multiple times to create several new bootstrap samples. By passing each sample through the same analysis we can then determine the different the effect random chance has on our results. Bootstrap can be performed at any point in the pipeline but applying it before machine learning would lead to us needing to perform the whole LOOCV machine learning 1000 times, once for each bootstrap sample. While this would have captured all uncertainty, it would have also biased the performance of the ML downwards by effectively reducing the sample size further for what is already a small study. In this study, we, therefore, chose to apply it to the output of a single LOOCV ML analysis of the original dataset. We, therefore, generate 1000 bootstrap samples on the predictions of each unique fold obtained by LOOCV, which are then sorted to determine the 95% quantiles, which correspond to the 95% confidence interval of the model’s results.

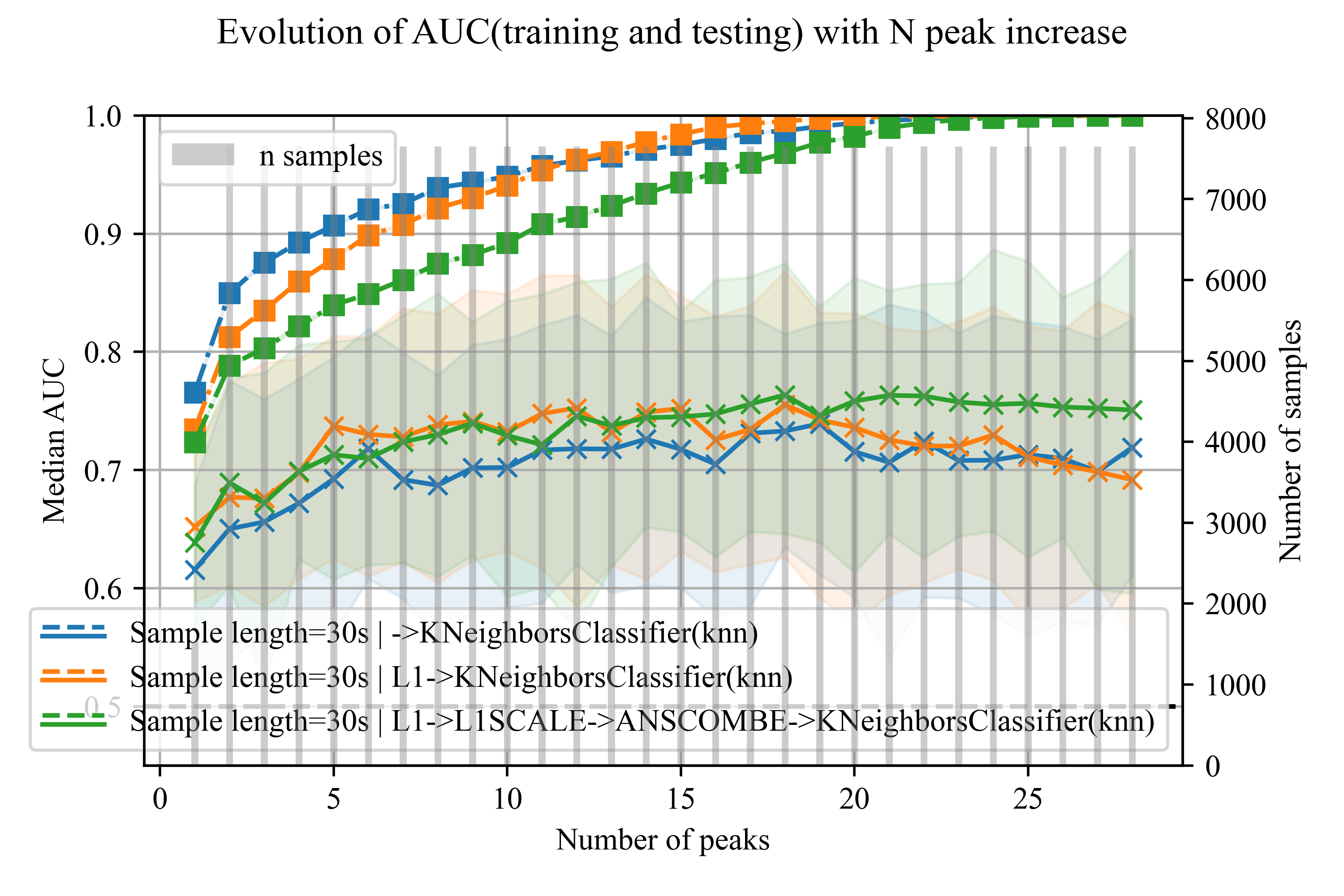

Figure S4 A.

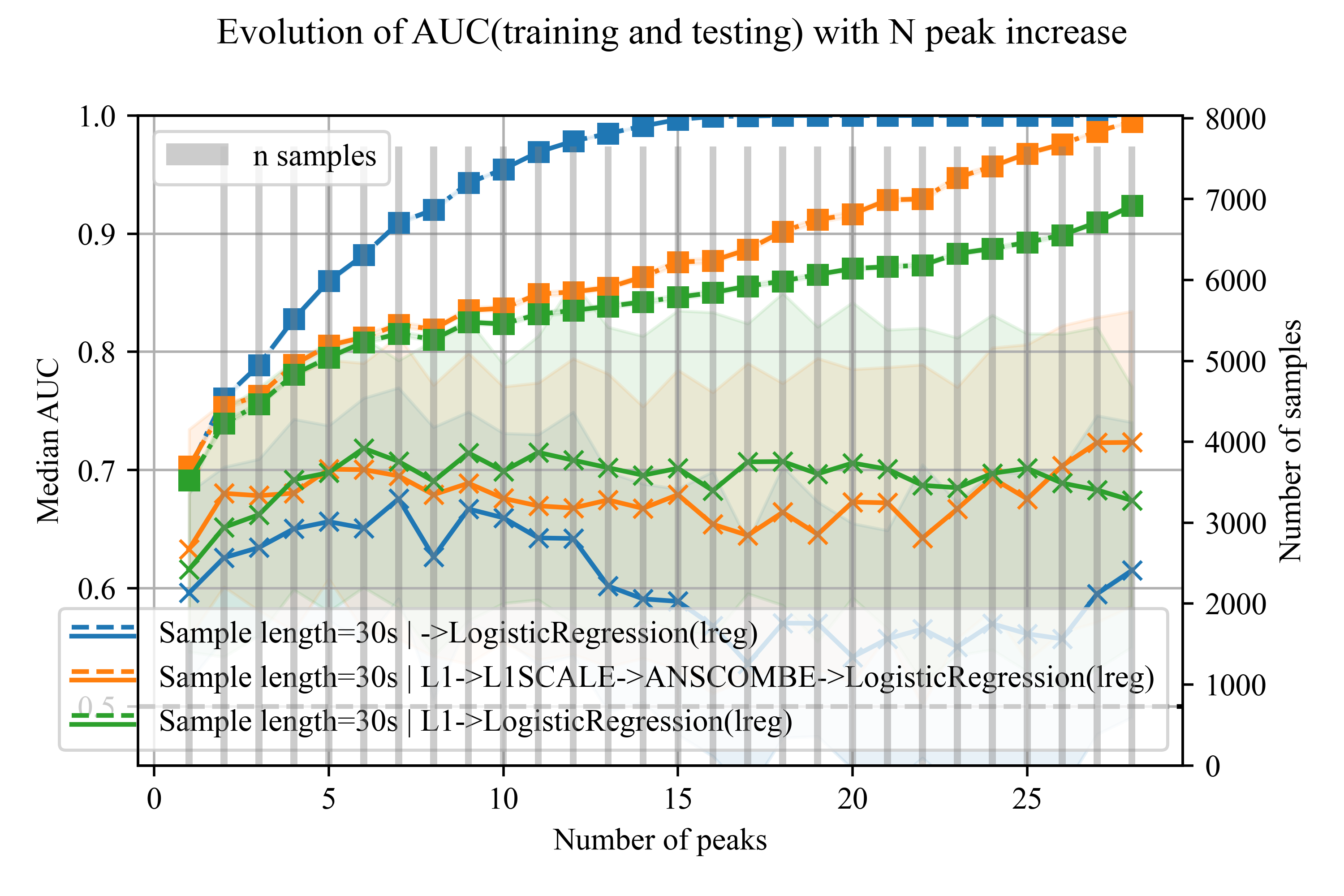

Figure S4 B.

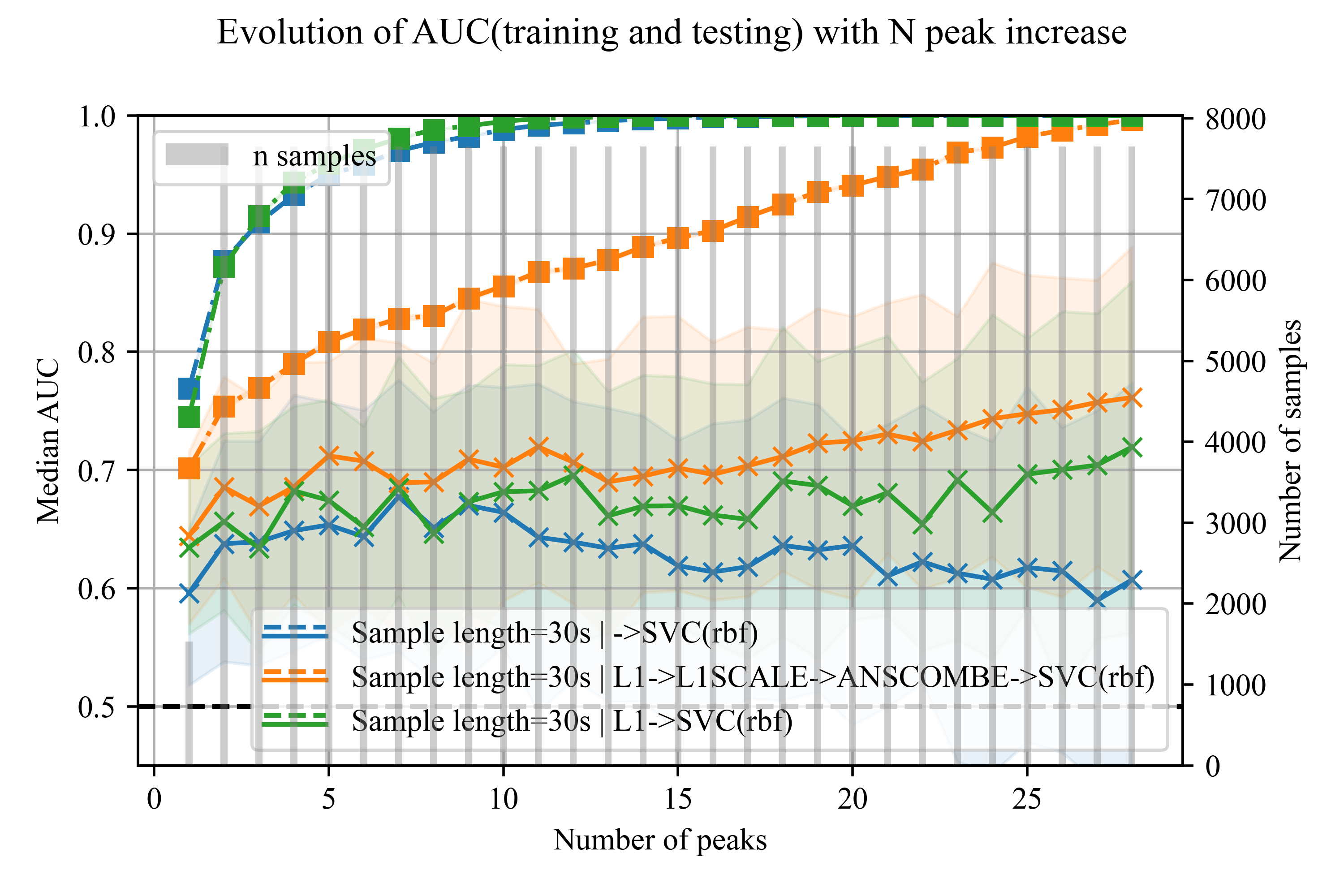

Figure S4 C.

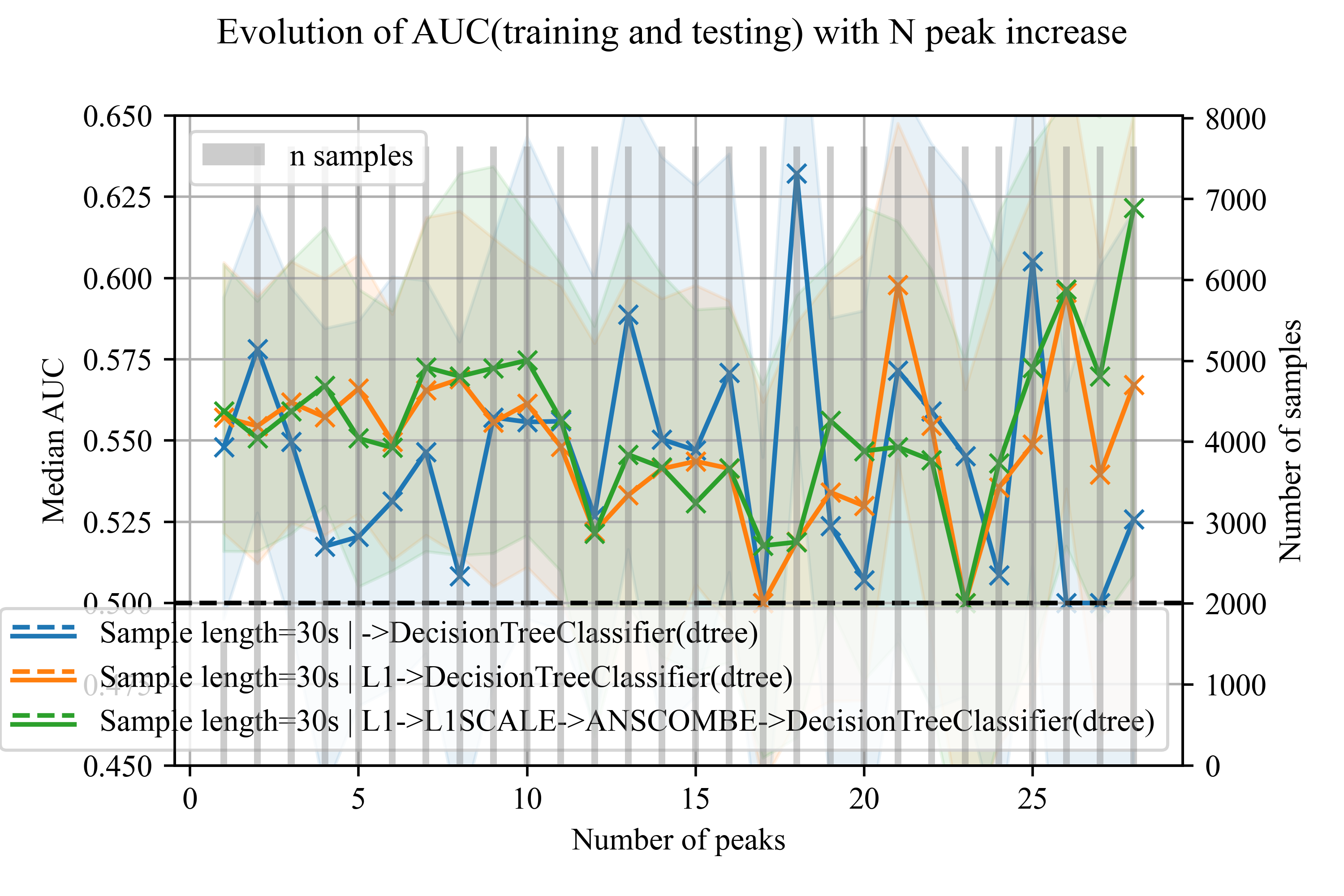

Figure S4 D.

**Supplementary  Table S2**

|  |  | *Actual* | |
| --- | --- | --- | --- |
|  |  | Unhealthy | Healthy |
| *Predicted* | Unhealthy | **True Positives** | False Positive |
|  | Healthy | False Negatives | **True Negative** |

Example of a confusion matrix. This example is based on our use case for the binary classification of healthy and unhealthy samples.

**Supplementary  Table S3**

| AUC testing | Class1 Precision testing | N training samples | N testing samples | N peaks | Sample length (seconds) | Classifier |
| --- | --- | --- | --- | --- | --- | --- |
| 0.75 (0.62-0.85) | 0.70 (0.51-0.83) | 7500 | 150 | 10 | 30 | SVC(rbf) |
| 0.75 (0.64-0.86) | 0.64 (0.50-0.79) | 7500 | 150 | 10 | 30 | SVC(rbf) |
| 0.71 (0.59-0.81) | 0.72 (0.58-0.84) | 7500 | 150 | 10 | 30 | SVC(rbf) |
| 0.72 (0.60-0.85) | 0.69 (0.52-0.84) | 7500 | 150 | 10 | 30 | KNeighborsClassifier(knn) |
| 0.70 (0.60-0.81) | 0.66 (0.50-0.82) | 7500 | 150 | 10 | 30 | LogisticRegression(lreg) |
| 0.69 (0.58-0.79) | 0.67 (0.51-0.81) | 7500 | 150 | 10 | 30 | LogisticRegression(lreg) |
| 0.67 (0.55-0.79) | 0.63 (0.45-0.76) | 7500 | 150 | 10 | 30 | SVC(rbf) |
| 0.65 (0.53-0.78) | 0.62 (0.43-0.78) | 7500 | 150 | 10 | 30 | SVC(rbf) |
| 0.54 (0.49-0.59) | 0.55 (0.39-0.68) | 7500 | 150 | 10 | 30 | DecisionTreeClassifier(dtree) |
| 0.55 (0.52-0.61) | 0.58 (0.41-0.72) | 7500 | 150 | 10 | 30 | DecisionTreeClassifier(dtree) |
| 0.63 (0.54-0.70) | 0.61 (0.48-0.75) | 7500 | 150 | 10 | 30 | LogisticRegression(lreg) |
| 0.54 (0.47-0.61) | 0.56 (0.41-0.72) | 7500 | 150 | 10 | 30 | DecisionTreeClassifier(dtree) |
| 0.66 (0.59-0.72) | 0.63 (0.44-0.74) | 1250 | 25 | 1 | 30 | SVC(rbf) |
| 0.65 (0.57-0.72) | 0.62 (0.48-0.73) | 1250 | 25 | 1 | 30 | SVC(rbf) |
| 0.65 (0.57-0.73) | 0.63 (0.52-0.76) | 1250 | 25 | 1 | 30 | KNeighborsClassifier(knn) |
| 0.65 (0.57-0.72) | 0.61 (0.46-0.75) | 1250 | 25 | 1 | 30 | KNeighborsClassifier(knn) |
| 0.61 (0.53-0.70) | 0.59 (0.45-0.73) | 1250 | 25 | 1 | 30 | LogisticRegression(lreg) |
| 0.62 (0.55-0.69) | 0.62 (0.49-0.77) | 1250 | 25 | 1 | 30 | LogisticRegression(lreg) |
| 0.64 (0.57-0.74) | 0.62 (0.48-0.78) | 1250 | 25 | 1 | 30 | LogisticRegression(lreg) |
| 0.60 (0.52-0.66) | 0.62 (0.43-0.73) | 1250 | 25 | 1 | 30 | SVC(rbf) |
| 0.61 (0.51-0.69) | 0.58 (0.47-0.70) | 1250 | 25 | 1 | 30 | KNeighborsClassifier(knn) |
| 0.53 (0.48-0.57) | 0.58 (0.43-0.71) | 1250 | 25 | 1 | 30 | DecisionTreeClassifier(dtree) |
| 0.53 (0.50-0.59) | 0.56 (0.43-0.69) | 1250 | 25 | 1 | 30 | DecisionTreeClassifier(dtree) |
| 0.53 (0.49-0.58) | 0.56 (0.42-0.69) | 1250 | 25 | 1 | 30 | DecisionTreeClassifier(dtree) |

**Predictive power of different classifier.** Shows the AUC and precision of different classifiers, SVC, KNN, LREG and DTREE for the dataset with and without augmentation.

**Supplementary  Figure legends**

Figure S1: **Cats age.** Boxplot showing the age of the cats used in this study.

Figure S2: **Cats accelerometer heatmap.** This heat map shows the accelerometery data of each cat during the study time. The raw data counts are pre-processed with the Anscombe and the Log transform. The excluded animals are highlighted in red.

Figure S3: **Schematic of Leave-one-out cross-validation.** In this example, there are 3 cats in the data-set, and each cat contains multiple peak samples 5.6. Leave one out cross-validation allows training (white blocks) on the data from all the cats but 1 which is excluded for testing (grey blocks), with this approach, a data set containing the data of 3 cats can be split 3 times.

Figure S4: **Evolution of the AUC with the increase of the number of peaks for different classifier**. While the x-axis shows the number of peaks used in the data set, the y-axis shows the Median AUC. The training AUC is shown by the dotted lines while the testing AUC is shown by the solid lines. All the datasets are fixed to 30 seconds sample, each subfigure is for the KNN (A), LREG (B), SVM (C) and DTREE (D) respectively.
