## Supplementary material for "Accelerometer-derived classifiers for early detection of degenerative joint disease in cats": Questionnaire Form

### Questionnaire 8: My 6 year old cat

Thank you for taking the time to update us about your cat's progress, now that he/she is 6 years old.

As before, this questionnaire contains a mixture of new questions, as well as some questions that you have answered previously – so that we can see what, if anything, has changed for your cat.

**Completing this questionnaire should be straightforward** and take about 20 minutes. If there are any questions which you do not wish to answer, please leave them blank and move to the next question.

Please return your completed questionnaire in the envelope enclosed.

Thank you for your help – information about the "Bristol Cats" and early results from the study will be available from our website.

**www.bristol.ac.uk/vetscience/cats**

FREEPOST RSHR-AGRJ-UABZ
Bristol Cats, Dr Jane Murray
University of Bristol
Langford House
Langford
BRISTOL
BS40 5DU
Tel/text: 07827 981412


### SECTION A: Your cat's household

**A1** Do you still have this cat?

| | Tick one box |
|---|---|
| Yes | |
| No | |

***If "Yes", please go to question A4***

**A2** Why do you no longer have this cat?

*Reason:*

**A3** Approximately how old was your cat when he/she left your household)?

Age of cat: …………….

***We are sorry to learn that you no longer have your cat. Please proceed to Section H to fill in your personal details.***

***If you would like to remain on our mailing list for newsletters please tick this box.*** ☐
***Thank you for participating in this study.***

**A4** How happy do you think your cat is?

| | Tick one box |
|---|---|
| Very happy | |
| Quite happy | |
| Not very happy | |
| Not at all happy | |
| I don't know | |

**A5** What factors contributed towards your answer above?

**A6** How many cats in total (including your 'Bristol cat(s)') currently live in this household?

………..

**A7**

| How frequently is this cat in a room where people smoke? | Tick one box |
|---|---|
| Never (currently and previously) | |
| Never (currently), but was previously (e.g. household member has stopped smoking) | |
| Less than once a week | |
| Once a week or more often | |

| | |
|---|---|
| A8 | Please indicate whether or not your cat has spent time during the last week in these types of places ***inside*** your home: |

| | ***Tick one box per row*** | | | |
|---|---|---|---|---|
| | Not available to my cat | Available to my cat | | |
| | | Used by cat | Not used by cat | Don't know if used by cat |
| A 'hiding' place that you have created for your cat (e.g. a 'pyramid' bed or cardboard box) | | | | |
| On a platform / ledge / raised area that your cat has previously used (but you didn't specifically create for your cat, e.g. on top of boiler, on windowsill) | | | | |
| On a platform / ledge / raised area that you have provided for your cat (e.g. a cat climbing frame bought from a shop) | | | | |

| | |
|---|---|
| A9 | Please indicate whether or not your cat has spent time during the last week in the following activities ***inside*** your home: |

| | ***Tick one box per row*** | | | |
|---|---|---|---|---|
| | Not available to my cat | Available to my cat | | |
| | | Used by cat | Not used by cat | Don't know if used by cat |
| Using a scratching post that you have provided for your cat | | | | |
| Using something else (other than a purpose built cat scratching post) to scratch on | | | | |
| Playing with bought or home-made cat toys | | | | |
| Playing with objects (not designed as a toy!) that the cat finds (e.g. a leaf) | | | | |

#### SECTION B: About your cat's activity levels and indoor/outdoor lifestyle

**B1** How often, if at all, do you believe that your cat

a) hunts outside for prey ?
b) eats prey that he/she has caught?

| | ***Tick one box per row*** | | | | |
|---|---|---|---|---|---|
| | Most days | Quite often (1-2 times/week) | Not very often (1-3 times/month) | Never | Don't know |
| Hunts outside for prey | | | | | |
| Eats prey that he/she has caught | | | | | |

**B2** Which of these statements best describes your cat's indoor/outdoor access?

| | *Tick one box* |
|---|---|
| Inside only – cat is not allowed outside | |
| Inside only – cat only goes out into enclosed 'run' or on a lead | |
| Inside and outside | |
| Outside only – cat is not allowed in the house | |

***If 'inside only', please go to section C.***
***If 'inside and outside', please continue with question B3.***
***If 'outside only', please go to question B4.***

**B3** Which of these statements best describes how much time your cat currently spends outside, **when he/she has unrestricted access to the outside**?

| | ***Tick one box*** |
|---|---|
| He/she hardly ever spends time outside | |
| He/she spends a little time outside, but most of his/her time is spent inside | |
| He/she spends roughly equal amounts of time inside and outside | |
| He/she spends a little time inside, but most of his/her time is spent outside | |
| He/she hardly ever spends time inside | |

**B4** To what extent does your cat have access to outdoor space beyond your garden?

| | ***Tick one box*** |
|---|---|
| Restricted at all times by a lead | |
| Restricted to the garden by a "cat proof" fence | |
| No restrictions | |
| Other *(please specify):* ………………………………………………………….. | |

### SECTION C: About your cat’s diet

***Many of these questions will be familiar as we would like to find out about any changes in your cat’s diet, appetite and food preferences.***

**C1** Which of the following sources of water do you know, or think, that your ‘Bristol cat’ drinks from?

| | ***Tick all that apply*** |
|---|---|
| Bowl of unfiltered tap water | |
| Bowl of filtered tap water | |
| Bowl of mineral water | |
| Cat drinking fountain – filtered tap water | |
| Cat drinking fountain – unfiltered tap water | |
| Toilet | |
| Running tap | |
| Outside source (stream, pond, puddles, etc) | |

**C2** How much do you feed your cat the following types of food?

| | ***Tick one box per row*** | | | | |
|---|---|---|---|---|---|
| | *Only food in diet* | *Major part (half or more) of daily diet* | *Minor part (less than half) of daily diet* | *Only feed occasionally* | *Never feed* |
| Commercial wet **adult** cat food (e.g. tins, pouches, foil packs) | | | | | |
| Commercial **adult** dry food (e.g. biscuits, kibbles) | | | | | |
| Uncooked/raw fresh food (e.g. fish, chicken) | | | | | |
| Cooked fresh food (e.g. fish, chicken) | | | | | |
| Cow’s milk/cream | | | | | |
| Cat milk | | | | | |

| | |
|---|---|
| C3 | Now, using the label on your cat's commercial wet food (e.g. tins, pouches, foil packs), please estimate which of these phrases best describes the total weight of commercial wet food that you think your cat eats each day. |

| | ***Tick one box*** |
|---|---|
| None | |
| 100g (e.g. one pouch or quarter of a standard-sized tin) | |
| 150g (e.g. one and a half pouches, or just over a third of a standard-sized tin) | |
| 200g (e.g. two pouches or half a standard-sized tin) | |
| 250g (e.g. two and a half pouches, or nearly two-thirds of a standard-sized tin) | |
| 300g (e.g. three pouches, or three-quarters of a standard-sized tin) | |
| Other *(please specify):* | |

| | |
|---|---|
| C4 | Please can you estimate (preferably using your kitchen scales) the weight (in grams) of dry food (e.g. biscuits, kibbles) that you think your cat eats each day, and enter this amount below. *(If your cat does not have dry food, please enter "0".)*<br>……………grams<br>Is this an accurate or estimated weight? |

| |
|---|
| Accurate |
| Estimate |

| | |
|---|---|
| C5 | If your cat receives home-prepared food (e.g. fresh fish), please describe what you would typically feed per day:<br><br>Please include information on the type (i.e. cut of meat, fish), preparation method and amount. |
| C6 | If you feed your cat commercial cat food and you mainly feed one or two brands (e.g Whiskas Tasty Textures / Felix Sensations etc, dry or tinned), what brands and varieties are they? |

| |
|---|
| Brand:<br>Variety: |
| Brand:<br>Variety: |

**C7**

If you use ‘wet food’, which of the following do you usually feed?

| | ***Tick all that apply*** |
| --- | --- |
| N/A, do not feed wet food | |
| Tins | |
| Pouches/sachets | |
| It varies | |
| Meat/fish in gravy | |
| Meat/fish in jelly | |
| Meat/fish pate | |
| Meat/fish Supermeat / Meaty Loaf | |
| Other (please specify): | |

**C8**

Using the following instructions, please assess the body condition of your ‘Bristol Cat’ and indicate the body condition score below:

To work out your cat’s individual body condition score, you need to do three checks:

1. Rib Check: Run both your hands, palms facedown across your cat’s ribcage on either side
2. Profile Check: View your standing cat from a side-on angle, this is best done if you are level with your pet
3. Overhead Check: Look down at your standing cat from an overhead angle

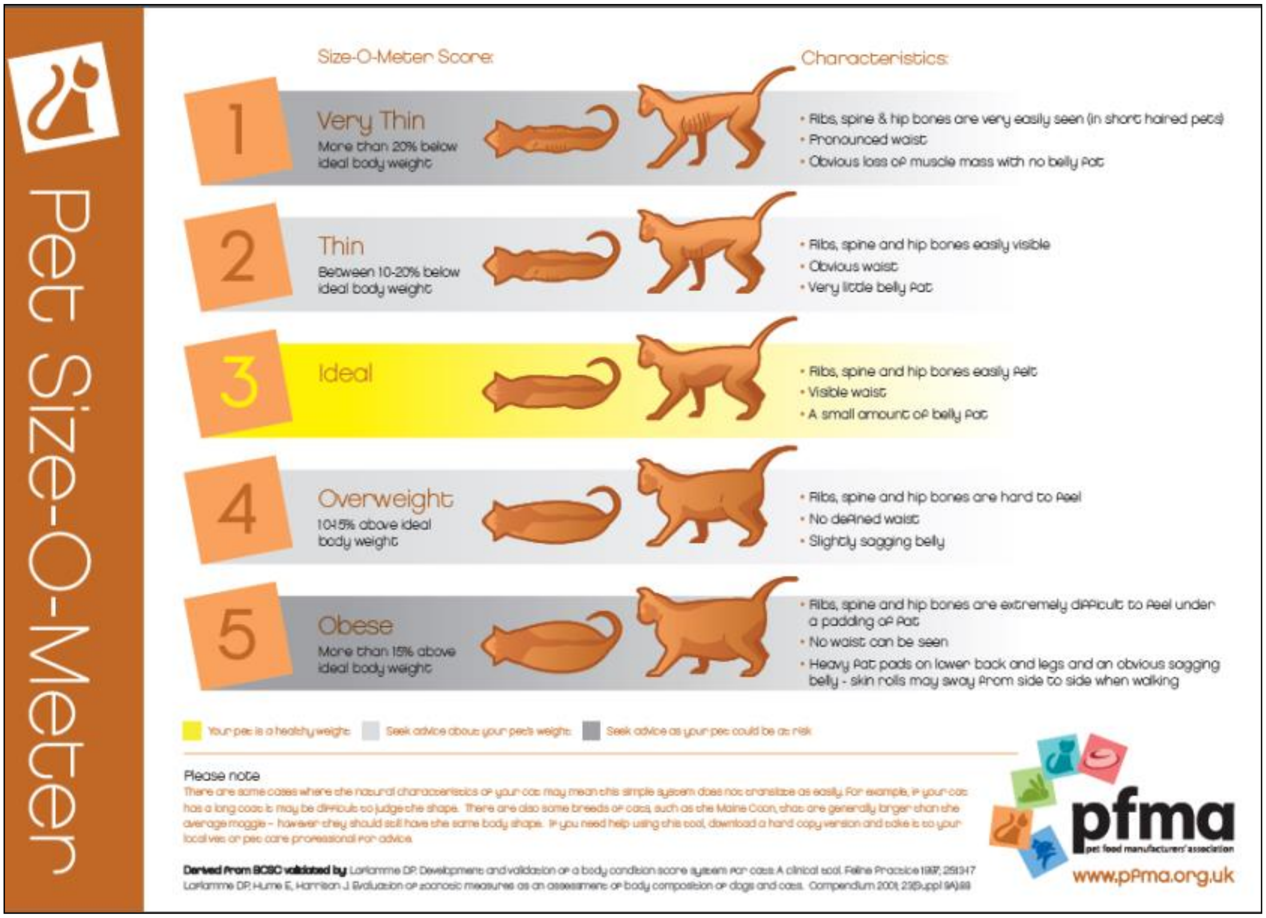

Source: www.pfma.org.uk/_assets/images/general/file/PFMA%20Cat%20PSOM%20Final%20Web%20Version%20070809.pdf

**Body condition score of ‘Bristol Cat’: ……….**

#### SECTION D: About your cat's health and veterinary contact

**D1** Has your cat visited a veterinary practice during the last ***12 months***?

| | ***Tick one box*** |
|---|---|
| Yes | |
| No | |

***If 'No', go to D5.***

**D2** What was the purpose of this visit / these visits?

| | ***Tick all that apply*** |
|---|---|
| Routine general health check | |
| Slimming club / weight watchers clinic | |
| Dental check | |
| Behavioural advice | |
| Vaccination | |
| Microchipping | |
| Neutering | |
| Flea prevention | |
| To treat fleas | |
| Worm prevention | |
| To treat worms | |
| Abscess / cat bite | |
| Attacked by dog | |
| Cat flu | |
| Coughing / wheezing | |
| Ear problem (e.g. ear mites) | |
| Urinary problem (e.g. cystitis, blocked bladder) | |
| Skin problem (e.g. itchy, excessive grooming, hair loss) | |
| Eye problem (e.g. conjunctivitis) | |
| Dental / tooth / mouth problem | |
| Lameness / limb problem, including broken or dislocated bone | |
| Checking / investigating heart problem (e.g. murmur) | |
| Vomiting / sickness | |
| Diarrhoea | |
| Cat 'off colour' | |
| Reduced appetite | |
| Increased appetite | |
| Increased thirst | |
| Weight loss | |
| Other *(please give full details):* | |

| | |
|---|---|
| D3 | Please use this space to provide further details of problems that have led to veterinary visits indicated above. If a vet has made a diagnosis please include this information. |
| D4 | Has your cat had any of the following diagnosed by a vet? |
| D5 | Please provide details of any medication (excluding routine flea/worming treatment) that your cat has received during the last 12 months. Information we are interested in includes: name of medication, dose, frequency medication given, date started, date finished (or length of course). |

| | ***Tick all that apply*** |
|---|---|
| Hyperthyroidism | |
| Heart disease | |
| Renal failure | |
| Diabetes | |
| Cancer | |
| None of these | |

| | |
|---|---|
| D6 | During the ***last 12 months,*** has your cat had any of the following illnesses/injuries/conditions which you felt were not serious enough to seek veterinary attention for? |

| | ***Tick one box per row*** | | |
|---|---|---|---|
| | *Yes* | *No* | *Not sure* |
| Fleas | | | |
| Worms | | | |
| Abscess / cat bite | | | |
| Attacked by dog | | | |
| Cat flu | | | |
| Coughing / wheezing | | | |
| Scratching his/her ears and/or shaking his/her head | | | |
| Urinary problem (e.g. cystitis, blocked bladder) | | | |
| Skin problem (e.g. itchy, excessive grooming, hair loss) | | | |
| Eye problem (e.g. conjunctivitis) | | | |
| Dental / tooth / mouth problem | | | |
| Lameness / limb problem | | | |
| Heart problem (e.g. previously detected murmur) | | | |
| Vomiting/sickness | | | |
| Diarrhoea | | | |
| Cat 'off colour' | | | |
| Reduced appetite | | | |
| Increased appetite | | | |
| Increased thirst | | | |
| Weight loss | | | |
| Other *(please specify):* | | | |

| | |
|---|---|
| D7 | Please use this space to provide further details of problems that your cat has had but which have not led to veterinary attention, as indicated above. |
| D8 | During the ***last 12 months***, have you seen any worms, fleas / flea dirt or signs suggestive of worms or fleas (e.g. scratching, worms in vomit or faeces)? |

| | ***Tick one box per row*** | | |
|---|---|---|---|
| | *Yes* | *No* | *Not sure* |
| Evidence of worms | | | |
| Evidence of fleas | | | |

| | |
|---|---|
| D9 | How many times, if at all, have you wormed or used flea treatment/prevention on your cat during the last 12 months? |

| | ***Tick one box per row*** | | | |
|---|---|---|---|---|
| | *Never* | *Once or twice* | *Three or more times* | *Don't know* |
| Wormer | | | | |
| Flea treatment | | | | |

| | |
|---|---|
| D10 | Is your cat insured? |

| | *Tick one box* | |
|---|---|---|
| | *Yes* | *No* |
| Insured | | |

| | |
|---|---|
| D11 | Please indicate the date of your cat's last vaccination. *(If not sure, please enter approximate month and year in space below)*: |

Date of last vaccination………/………/………..

Or, approximate date:……….…..(month) ……..…….(year)

☐ Not applicable: never been vaccinated (**go to D15**)

| | |
|---|---|
| D12 | Was your cat vaccinated at the practice that was your 'usual' veterinary practice at that time? |

| | ***Tick one box*** |
|---|---|
| Yes – usual veterinary practice (please also tick if you are a vet and usually vaccinate your own cat) | |
| No – I went to a different practice on this occasion | |

| | |
|---|---|
| D13 | Are the details of your cat's most recent vaccination recorded on his/her vaccination card? |

| | ***Tick one box*** |
|---|---|
| Yes | |
| No | |
| Can't remember | |

| | |
|---|---|
| D14 | At the last vaccination, which of the following diseases did your vet recommend your cat was vaccinated against and which diseases was your cat actually vaccinated against? |

| | ***Tick all that apply*** | | | | | |
|---|---|---|---|---|---|---|
| | ***Vet recommended*** | | | ***Cat vaccinated against*** | | |
| ***Disease*** | ***Yes*** | ***No*** | ***Not sure*** | ***Yes*** | ***No*** | ***Not sure*** |
| Bordetella | | | | | | |
| Cat flu (Feline Herpes Virus (FHV-1) / Feline Calicivirus (FCV)) | | | | | | |
| Feline Infectious Enteritis (FIE) or Panleucopenia | | | | | | |
| Feline Leukaemia Virus (FeLV) | | | | | | |
| Feline Chlamydophilosis | | | | | | |
| Rabies | | | | | | |

| | |
|---|---|
| D15 | Excluding emergency/out of hours appointments, do you use different veterinary practices for different problems/treatments for this cat, or do you use the same veterinary practice for everything? |

| | ***Tick one box*** |
|---|---|
| Same practice for everything | |
| Different practices for different problems/illnesses/treatments | |
| Prefer not to answer this question | |
| Other *(please specify):* | |

| | |
|---|---|
| D16 | Is your 'Bristol cat' neutered (desexed)? |

| | ***Tick one box*** |
|---|---|
| Yes – at or before 5 years of age | |
| Yes – since 5 years of age | |
| No | |

| | |
|---|---|
| D17 | How frequently, if at all, do you do the following to help keep your cat's teeth and mouth healthy? |

| | ***Tick one box per row*** | | | | |
|---|---|---|---|---|---|
| | *Every day* | *A few times a week* | *Once a week* | *Less frequently* | *Never* |
| Brush teeth | | | | | |
| Use dental gel or mouth rinse | | | | | |
| Use food or water additive | | | | | |
| Feed dental treats | | | | | |
| Feed a special dental health diet | | | | | |
| Feed home-prepared fresh food | | | | | |
| Other *(please specify):* | | | | | |

| | |
|---|---|
| D18 | If you use any 'dental health' products mentioned in the question above, please name the product(s) used: |

| | |
|---|---|
| D19 | During the last 12 months, has a vet/vet nurse commented on the health of your cat's teeth and mouth? |

| | ***Please tick the comment that applies best*** |
|---|---|
| Yes – advised that teeth and mouth are in good health | |
| Yes – advised that cat has some dental/oral disease and that dental treatment (under anaesthetic) may be necessary in the future | |
| Yes – advised that that cat has a 'scale and polish' (under anaesthetic) | |
| Yes – advised that cat has dental/oral disease and recommended that dental treatment under anaesthetic (excluding a 'scale and polish only') was needed | |
| No – no comment on teeth/mouth made | |
| N/A – has not seen a vet or vet nurse in the past 12 months | |

| | |
|---|---|
| D20 | Has your cat had any dental work carried out during the last 12 months by the vet? |

| | ***Tick one box*** |
|---|---|
| Yes | |
| No | |

***If 'no', please go to question D22.***

| | |
|---|---|
| D21 | If your cat has had dental work carried out during the last 12 months, please provide further information below regarding the work carried out: |

| | ***Tick one box*** |
|---|---|
| Scale and polish only | |
| A few teeth extracted (e.g. 1 or 2) | |
| A moderate number of teeth extracted (e.g. 3-6) | |
| A lot of teeth extracted (e.g. 7 or more) | |
| Other (please specify, including reasons for extractions, if known:) | |

| | |
|---|---|
| D22 | During the last 12 months, have you seen your cat drinking, urinating or defecating? |

| | ***Tick one box for each row*** | |
|---|---|---|
| | *Yes* | *No* |
| Drinking | | |
| Urinating | | |
| Defecating | | |

| | |
|---|---|
| D23 | Please indicate whether you are aware of any changes in your cat in the areas of drinking and urination, during the last 12 months. |

| | ***Tick one box for each row*** | | | |
|---|---|---|---|---|
| | Not aware of any changes | No change | Increased | Decreased |
| *Amount of water that cat drinks* | | | | |
| *Amount of urine that cat passes* | | | | |

| | |
|---|---|
| D24 | If you think that your cat has been drinking more, or less, water during the last 12 months, please indicate the reason(s) for your answer. |

| | ***Tick all that apply*** |
|---|---|
| Water bowl needs refilling more/less frequently | |
| I see the cat drinking inside more/less often | |
| I see the cat drinking outside more/less often | |
| Other *(please specify)* | |

| | |
|---|---|
| D25 | Which, if any, of the following have you been aware of whilst watching your cat urinating? |

| | ***Tick one box for each row*** | | |
|---|---|---|---|
| | *Yes* | *No* | *N/A have not seen cat urinating* |
| He/she strains or appears to have difficulty urinating | | | |
| He/she has passed blood when urinating | | | |
| He/she vocalises (e.g. miaows) before or during urination | | | |
| He/she sometimes urinates in different locations (around the house and/or outside) | | | |

D26 Using the diagrams below, please indicate how frequently during the last week, if at all, has your cat been grooming, scratching, biting, licking, chewing, nibbling, rubbing (out of discomfort rather than 'normal rubbing or grooming behaviour') him/herself in <u>any area</u> within each of these marked regions.

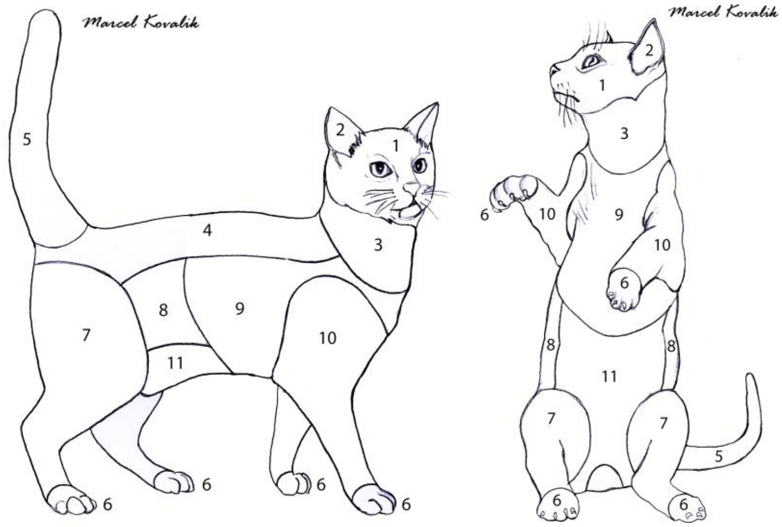

| | *Tick one box for each row* | | | | |
|---|---|---|---|---|---|
| | Almost continuously | A lot of the time and/or for long spells of time (including when eating, playing or being distracted) | A moderate amount of time (but not when eating, playing or being distracted) | Only occasionally (and out of discomfort rather than 'normal rubbing or grooming behaviour') | Not at all (out of discomfort rather than 'normal rubbing or grooming behaviour') |
| 1 (head) | | | | | |
| 2 (ears) | | | | | |
| 3 (neck) | | | | | |
| 4 (back and base of tail) | | | | | |
| 5 (tail) | | | | | |
| 6 (paws) | | | | | |
| 7 (back legs and thigh, excluding paws) | | | | | |
| 8 (flank) | | | | | |
| 9 (chest and sides) | | | | | |
| 10 (front legs and shoulders, excluding paws) | | | | | |
| 11 (tummy) | | | | | |

D27 Using the same two diagrams, please indicate which of the following list, if any, you have noticed on your cat:

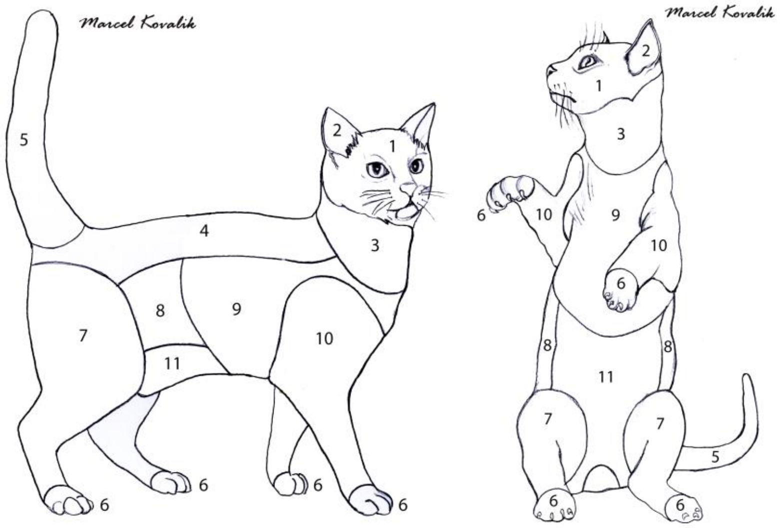

| | *Tick all that apply one box* | | | | | | |
|---|---|---|---|---|---|---|---|
| | Bald patches / clumps of hair missing / falling out | General thinning of the coat (including short barbered hair but excluding usual moulting) | Scabs / crusts | Lumps / bumps / swellings | Redness of the skin | Bleeding | None of these - hair and skin appear normal |
| 1 (head) | | | | | | | |
| 2 (ears) | | | | | | | |
| 3 (neck) | | | | | | | |
| 4 (back and base of tail) | | | | | | | |
| 5 (tail) | | | | | | | |
| 6 (paws) | | | | | | | |
| 7 (back legs and thigh, excluding paws) | | | | | | | |
| 8 (flank) | | | | | | | |
| 9 (chest and sides) | | | | | | | |
| 10 (front legs and shoulders, excluding paws) | | | | | | | |
| 11 (tummy) | | | | | | | |

#### SECTION E: What is normal for your cat at the moment?

E1 During a 'typical' week, which of these phrases best describes the ***usual*** consistency of your cat's faeces?

| | ***Tick one box*** |
|---|---|
| Have not seen faeces | |
| Dry/hard | |
| Firm | |
| Soft/loose | |
| Runny/watery | |
| Varies | |

E2 Please select the statement that best applies to your cat for each of these activities.

| | ***Tick one box per row*** | | | |
|---|---|---|---|---|
| My cat…. | No | Maybe | Yes | N/A |
| is less willing to jump up or down than he/she was 18 months ago | | | | |
| will only jump up or down from lower heights | | | | |
| shows signs of being stiff at times | | | | |
| is less agile than he/she was 18 months ago | | | | |
| shows signs of lameness or limping | | | | |
| has difficulty getting in or out of the cat flap | | | | |
| has difficulty going up or down stairs | | | | |
| cries when picked up | | | | |
| has accidents outside the litter tray | | | | |
| spends less time grooming than he/she did 18 months ago | | | | |
| is more reluctant to interact with me than he/she was 18 months ago | | | | |
| plays less (e.g. with other animals and/or toys) than he/she did 18 months ago | | | | |
| sleeps more and/or is less active than he/she did 18 months ago | | | | |
| cries out loudly for no apparent reason | | | | |
| appears forgetful or disorientated | | | | |

E3 Please rate how well you think your cat is able to carry out the following activities during a 'typical' week?

| | ***Tick one box per row*** | | | |
|---|---|---|---|---|
| | Very well | Well | Adequately | Not well |
| Grooming | | | | |
| Eating | | | | |
| Moving around | | | | |

| | |
|---|---|
| E4 | Please indicate how easily your cat can jump up onto the following places: |

| | Tick one box per row | | | | | |
|---|---|---|---|---|---|---|
| | Very easily | Quite easily | Not very easily | With a lot of difficulty | Cannot | Don't know |
| Sofa | | | | | | |
| Your bed | | | | | | |
| Kitchen work surface | | | | | | |
| Kitchen / dining table | | | | | | |

We are assuming the following *approximate* standard heights.

Sofa: 40 cm (16")  Work surface: 90 cm (36")
Bed: 55 cm (22")  Table: 78 cm (31")

If your furniture is a 'non-standard' height, please provide a measurement at the end of the questionnaire in the space for further information.

| | |
|---|---|
| E5 | Please rate your perception of your cat's activity levels during the past week using the options below. |

| | ***Tick one box*** |
|---|---|
| Very active | |
| Quite active | |
| Not very active | |
| Not at all active | |

| | |
|---|---|
| E6 | Do you believe that there are currently any 'external' factors that are affecting your cat's mental or physical wellbeing? (E.g. bullying from another cat, moving house). |

| | ***Tick one box*** |
|---|---|
| Yes | |
| No | |

***If 'No', go to E8.***

| | |
|---|---|
| E7 | We would be grateful if you could provide further details about these 'external' factors. |
| E8 | Please rate your perception of your cat's overall quality of life ***during the past week*** using the options below. |

| | ***Tick one box*** |
|---|---|
| Excellent | |
| Good | |
| Average | |
| Fair | |
| Poor | |

| | |
|---|---|
| E9 | What factors contributed towards your selection of this rating? |

### SECTION F: About your cat's behaviour and characteristics

F1

***If you only have one cat, please go to F2.***

If you have more than one cat, which of these statements best describes how your 'Bristol Study cat' interacts with other cats in the household? He/she…

| | ***Tick one box per row*** | |
|---|---|---|
| | *Yes* | *No* |
| Sleeps in the same room as another cat, but not close together | | |
| Shares a sleeping place with another cat | | |
| Sleeps with another cat where they are touching each other | | |
| Grooms another cat | | |
| Is groomed by another cat | | |
| Rubs on another cat | | |
| Is rubbed on by another cat | | |
| Chases another cat | | |
| Is chased by another cat | | |
| Plays with another cat | | |
| Hisses or spits at another cat | | |
| Is hissed or spat at by another cat | | |
| Is reluctant to pass another cat in a narrow space (e.g. doorway) | | |
| Blocks or inhibits the movement of another cat | | |

F2

*Does your cat display any undesirable behaviours?*

| | ***Tick one box*** |
|---|---|
| Yes | |
| No | |

***If 'No', please go to section G.***

F3

If yes, please describe this/these behaviour(s).

Behaviour 1:

Behaviour 2:

Behaviour 3:

Other undesirable behaviours:

F4

For each of the behaviours listed above, (or for the first three listed), please provide the following information:

| Behaviour | Is this behaviour a problem to you? *Tick all that apply* | | Please indicate whether or not you have sought help for these behaviour problems *Tick all that apply* | |
|---|---|---|---|---|
| | Yes | No | Yes | No |
| Behaviour 1 | | | | |
| Behaviour 2 | | | | |
| Behaviour 3 | | | | |

***If 'No' to all behaviours, please go to section G.***

F5

Please tick which of these sources you have sought help from for each of these behaviour problems, (or for the first three listed).

| | *Tick all that apply* Behaviour 1 | Behaviour 2 | Behaviour 3 |
|---|---|---|---|
| Vet | | | |
| Vet behaviourist | | | |
| Behaviourist | | | |
| Vet Nurse | | | |
| Other members of vet practice staff | | | |
| Friend | | | |
| Books | | | |
| Animal welfare organisation staff/online resources | | | |
| Online sources not mentioned above (including forums) | | | |
| Other *(please provide details)* | | | |
| Have not sought help | | | |

### SECTION G: Your cat's neighbourhood

**G1** If your cat has been registered with a veterinary practice (for the first time, or with a different veterinary practice), since you completed the last questionnaire for the 'Bristol Cats' study, and you are happy for us to access your cat's veterinary records, please provide the **name and address** of your cat's new veterinary practice below:

Name of veterinary practice:

Address:

**G2** Have you moved house within the last 12 months?

| | Tick one box |
|---|---|
| Yes | |
| No | |

***If 'No', please go to G5.***

**G3** Which of these phrases best describes the location of your home?

| | Tick one box |
|---|---|
| In a rural area | |
| In a village or suburban location | |
| In a town or city location | |
| Don't know | |

**G4** Do you have a garden? (Include communal gardens.)

| | Tick one box |
|---|---|
| Yes | |
| No | |

***The next few questions check for the contact your cat has with local dogs and cats, possibly due to new dogs/cats moving to your area, or because you have moved house.***

**G5** How many dogs (from other households) do you know of that are in your immediate neighbourhood? (I.e. dogs that your cat might see and/or hear outside regularly.)

| | Tick one box |
|---|---|
| None | |
| 1-5 | |
| 6-10 | |
| 11 or more | |

**G6** How many cats (from other households) do you know of that are in your immediate neighbourhood? (I.e. cats from other household that your cat might see and/or hear outside regularly.)

| | Tick one box |
|---|---|
| None | |
| 1-5 | |
| 6-10 | |
| 11 or more | |

***If 'None', go to G12.***

G7 Do any of these cats stare into your house through catflaps, doors or windows?

| | ***Tick one box*** |
|---|---|
| Yes | |
| No | |
| Don't know | |

G8 In which of these ways does your cat react if he/she sees any of these cats in his/her house or garden?

| | ***Tick all that apply*** |
|---|---|
| Ignores them | |
| Stays still | |
| Hisses or spits | |
| Chases them | |
| Swipes his / her paw | |
| Runs away | |
| Rubs against them | |
| Licks or grooms them | |
| Plays with them | |
| Skirts around them | |

***If your cat is an 'indoor only' cat, please go to G12.***

G9 How many cats (from other households) does your cat come into contact with at least once a week, in each of the following categories?

| | ***Tick one box per row*** | | | |
|---|---|---|---|---|
| | *None* | *1-3* | *4 or more* | *Don't know* |
| 'Friends' of my cat (e.g. they play, spend time together) | | | | |
| 'Acquaintances' of my cat (e.g. they meet occasionally but there is little contact between them) | | | | |
| 'Enemies' of my cat (e.g. they fight, one of them will chase the other, they avoid each other) | | | | |
| Other *(please specify):* | | | | |

G10 Do any of these cats come into your house?

| | ***Tick one box*** |
|---|---|
| Yes | |
| No | |
| Don't know | |

G11 Do any of these cats come into your garden?

| | ***Tick one box*** |
|---|---|
| Yes | |
| No | |
| Don't know | |

| | | | |
|---|---|---|---|
| G12 | During the last six months, how often, if at all, do you think your cat has been involved in a fight with another cat? | ***Tick one box per column*** | |
| | | *Cats within the household* | *Cats from another household* |
| | Not applicable (e.g. no other cats in household, cat does not have outside access) | | |
| | Never | | |
| | Once a month or less often | | |
| | 2-4 times a month | | |
| | Once a week or more often | | |

G13 Please use this space to tell us about any ***major changes*** relating to your cat's **environment / behaviour / diet / health** that have taken place over the last 12 months and which have not been covered in this questionnaire.

G14 All things considered, how willing would you be to take on the life your cat is now living?

| | ***Tick one box*** |
|---|---|
| Very willing | |
| Quite willing | |
| Not very willing | |
| Not at all willing | |
| I don't know | |

G15 Please use this space for any additional comments you would like to add:

### SECTION H: Final details

Finally, please complete the details below for our records. This final section is very important and enables us to link this questionnaire with others you have completed for your cat. Please be reassured that this information is strictly confidential and will be used for no other purposes. Your contact details will ONLY be used for the purposes of the 'Bristol Cats' study.

| |
|---|
| **Date of questionnaire completion** |
| **IDENTIFYING INFORMATION:** |
| **Bristol Cat 'Owner ID'** |
| **Bristol Cat 'Cat ID'** |
| **Name of cat** |
| **Your name** |
| **Address** |
| **Email address** |
| **Contact telephone number** |

***Thank you very much for your time and help in completing this questionnaire.***

***We really appreciate the time that you have taken to complete this questionnaire to tell us about your cat. The information you give us about your cat will help us to help cats in the future. If you have any questions, please contact a member of the study team.***

Signature: ……………………………..…………Date: ……………………….

Please return your completed questionnaire in the envelope enclosed.

Freepost RSHR-AGRJ-UABZ

Bristol Cats: Dr Jane Murray

University of Bristol, Langford House, Langford, BRISTOL, BS40 5DU
